# De-novo DNA Methylation of Bivalent Promoters Induces Gene Activation through PRC2 Displacement

**DOI:** 10.1101/2025.02.07.636872

**Authors:** Henriette O’Geen, Alina Mihalovits, Benjamin D. Brophy, He Yang, Maxwell W. Miller, Chelsey J. Lee, David J. Segal, Marketa Tomkova

## Abstract

Promoter DNA methylation is a key epigenetic mark, commonly associated with gene silencing. However, we noticed that a positive association between promoter DNA methylation and expression is surprisingly common in cancer. Here, we use hit-and-run CRISPR/dCas9 epigenome editing to evaluate how deposition of DNA methylation can regulate gene expression dependent on pre-existing chromatin environment. While the predominant effect of DNA methylation in non-bivalent promoters is gene repression, we show that in bivalent promoters this often leads to gene activation. We demonstrate that gain of DNA methylation leads to reduced MTF2 binding and eviction of H3K27me3, a repressive mark that guards bivalent genes against activation. Our cancer patient data analyses reveal that in cancer, this mechanism likely leads to activation of a large group of transcription factors regulating pluripotency, apoptosis, and senescence signalling. In conclusion, our study uncovers an activating role of DNA methylation in bivalent promoters, with broad implications for cancer and development.

## Introduction

DNA methylation (5-methylcytosine, 5mC) is a major epigenetic modification that plays important roles in the regulation of gene expression, silencing of transposons, X chromosome inactivation, genomic imprinting, genomic stability, and mammalian development^1,2^. In the 1970s, it was proposed that DNA methylation might be responsible for regulation and maintenance of stable gene expression across cell division^3,4^. Since then, it has been shown that DNA methylation can directly cause gene repression^5,6^ and numerous studies have further established the link between promoter DNA methylation and gene silencing^7–10^. This repressive role is thought to act through recruitment of histone deacetylases, promoting and stabilising repressive heterochromatin formation by 5mC-binding proteins, blocking binding of activating transcription factors (TFs), or inducing RNA polymerase II stalling^7,8,11^. Next to a direct causal role in gene silencing, DNA methylation can serve also as a ‘lock’ to reinforce a previously silenced state^7,12^. While DNA methylation is sometimes associated with active transcription^13,14^, especially when located in gene bodies^7,15^, it is broadly accepted that DNA methylation in promoters is a key repressive mark^1,7–10,12,16^.

In cancer, aberrant 5mC distribution has been observed across most cancer types, including promoter hypermethylation of tumour suppressors, hypomethylation of oncogenes, and global hypomethylation linked to genomic instability^1^. Epigenetic aberrations have emerged as having a pivotal role in the development of cancer: epigenetic abnormalities are able to give rise to every hallmark of cancer, and epigenetics is increasingly being used in cancer diagnostics and therapeutics^17–20^.

Interestingly, a fraction of promoters marked with both repressive H3K27me3 and activating H3K4me3 histone marks in embryonic stem cells (ESCs) are particularly prone to hypermethylation in cancer^21–24^. These so-called *bivalent* domains are typically located near developmental genes and transcription factors (TFs), and maintain genes in a poised state, prepared to be activated or permanently repressed in different lineages during differentiation^25–28^. It has been thought that replacement of histone modifications by DNA methylation in cancer might serve to ensure more stable gene repression^7,21,29–32^. However, more recent large scale high-throughput sequencing data revealed that genes associated with bivalent chromatin in ESCs are in fact often upregulated in cancer^33^.

In this study, we sought to investigate this discrepancy and uncover the relationship between DNA methylation and transcription with respect to bivalent chromatin. Using a combination of cancer patient data analysis and CRISPR/dCas9 hit-and-run epigenome editing, we show a novel role of promoter DNA methylation in activation of bivalent genes. We see long-lasting DNA methylation gain in bivalent promoters deposited by epigenome editing, which leads to decreased H3K27me3 and increased RNA expression. This mechanism provides a likely explanation as to why hundreds of genes bivalently marked in normal tissue become hypermethylated and activated in cancer patients. These genes comprise dozens of developmental transcription factors that regulate numerous cancer pathways involved in apoptosis, cell cycle, senescence, and pluripotency signalling. In summary, while promoter DNA methylation acts primarily as a repressive mark outside bivalent chromatin, our results implicate an activating role of DNA methylation in the context of bivalent chromatin.

## Results

### Bivalent chromatin in normal tissue predisposes to DNA hypermethylation and gene activation in cancer

In order to quantify the relationship between promoter DNA methylation and gene expression in cancer, we first analysed DNA methylation HM450K array and gene expression RNA-seq data from 3,777 patients in The Cancer Genome Atlas (TCGA) across seven cancer types (Fig. 1a, Supplementary Table 1). Initially, we focused on all differentially expressed (DE) genes with differentially methylated (DM) CpGs in their promoter (DM-DE genes) (Fig. 1b, see Methods). To avoid signal due to gene-body driven effects^7,15^, only CpGs in promoter regions up to 1,500 bp upstream of transcription start sites (TSS) were considered in this analysis.

**Fig. 1:**
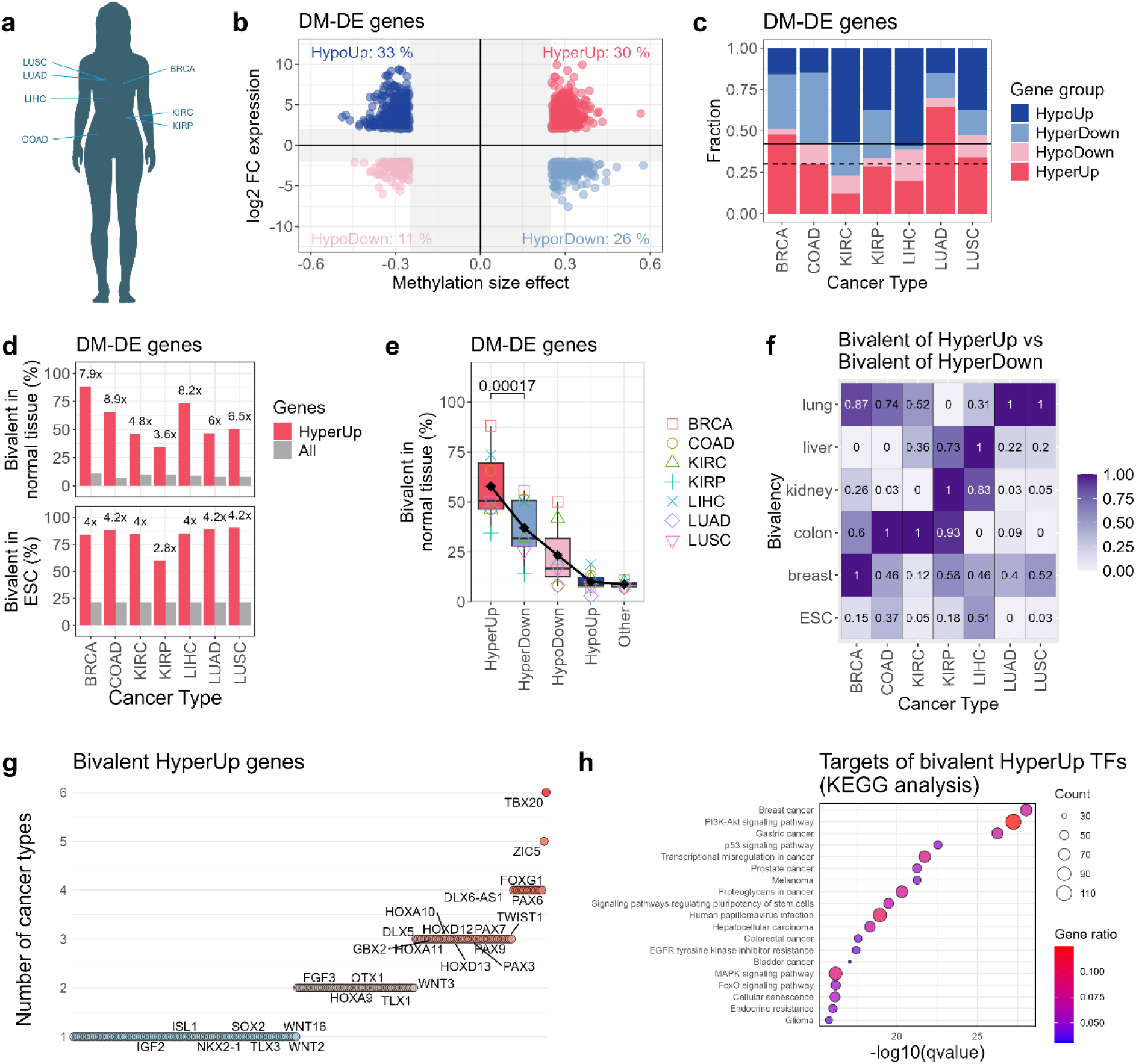
Hypermethylated and upregulated genes are frequent in cancer and largely explained by bivalent chromatin in matched normal tissue. **a**, DNA methylation (EPIC array) and gene expression (RNA-seq) data from 7 TCGA cancer types were analysed: BRCA = breast invasive carcinoma, COAD = colon adenocarcinoma, KIRC = kidney renal clear cell carcinoma, KIRP = kidney renal papillary cell carcinoma, LUSC = lung squamous cell carcinoma, LUAD = lung adenocarcinoma, LIHC = liver hepatocellular carcinoma. **b,** All genes that are differentially methylated (DM) and differentially expressed (DE) in at least one cancer type. The x-axis shows methylation size effect (hypermethylated in cancer on the right), computed as average cancer minus normal CpG methylation of the differentially methylated promoter CpGs of that gene. The y-axis shows expression size effect (upregulated in cancer on the top), computed as log2 fold-change of expression in cancer/normal. For genes that are significantly differentially methylated and expressed in more than one cancer type, the average value across the significant cancer types is shown (genes that fall in different quadrants in different cancer types have been excluded). **c,** Fraction of genes in each quadrant in (b), shown for each cancer type. The dashed line represents the average of HyperUp groups across cancer types (30%). The solid line represents the average of HyperUp plus HypoDown groups (42%). **d,** Percentage of H1 embryonic stem cell (ESC) line (bottom) and normal tissue (top) bivalent genes in HyperUp genes (red) vs all genes (grey). The numbers above the red bars represent fold change between the red and grey bars. **e,** Percentage of genes in each quadrant of (a) that are bivalent (marked with H3K4me3 and H3K27me3 in the promoter) in matched normal tissue (data from ENCODE). P-value was determined by two-sided paired T-test. **f,** Enrichment of promoters marked as bivalent in different normal tissues (rows) in genes that are HyperUp in different cancer types (columns), computed as column-normalized (0 as minimum, 1 as maximum) value of (percentage of bivalent in HyperUp) / (percentage of bivalent in HyperDown). **g,** A scatterplot showing all bivalent HyperUp genes. The y-axis represents the number of cancer types where this gene is bivalent and hypermethylated and upregulated. Selected genes are labelled. **h,** KEGG pathway analysis of targets of bivalent HyperUp transcription factors (TFs), as identified by CollecTRI^41^. Gene ratio represents k/n, where n is the number of all the target genes (912) and k is the number of the target genes that participate in the given pathway. Count represents the value of k. The top pathways with lowest q value are shown in the plot. The full list of significantly enriched pathways is presented in Supplementary Table 4.

Given that promoter DNA methylation has, in general, been associated with gene repression, we would expect the DM-DE genes to fall mainly in the hypomethylated and upregulated (HypoUp) or hypermethylated and downregulated (HyperDown) categories. However, a *positive* association between differential methylation and differential expression is surprisingly frequent, comprising together a median of 42% of DM-DE genes across cancer types (Fig. 1c, Supplementary Figure 1). In particular, the category of hypermethylated and upregulated (HyperUp) genes is especially common, covering a median of 30% of DM-DE genes (Fig. 1c, Supplementary Table 2).

Next, we used histone mark measurements from ENCODE to characterize the group of HyperUp genes. On average, 85% of the HyperUp genes are explained by a bivalent combination of H3K27me3 and H3K4me3 marks in the gene promoters in H1 ESC (Fig. 1d). This is 4-fold higher than expected by chance (Fig. 1d).

Bivalent chromatin is most commonly studied in embryonic and pluripotent stem cells, where it is most prevalent and gradually diminishes during differentiation^25,31^. However, bivalent regions can be detected across a wide range of cell types and tissues. We were therefore interested in whether our results reflect a specific property of bivalency in ESC, or whether bivalent chromatin in pre-cancerous/normal tissue is responsible for the methylation and expression gain in cancer. Therefore, we repeated the analysis using normal tissue bivalent maps from ENCODE, used as a proxy of the pre-cancerous tissue-of-origin (see Methods). As expected, the percentage of bivalent genes was reduced in general (Fig. 1d-e). However, the enrichment of bivalent genes in the group of HyperUp genes was even stronger for bivalent maps from matched normal tissues (6.5-fold higher than expected by chance) than for bivalent maps from ESC cells (Figure 1d). In particular, these bivalent genes were significantly more enriched in HyperUp than HyperDown group (Fig. 1e). Moreover, when comparing this enrichment for bivalent maps from across the different tissues, the tissue-matched bivalent maps showed overall the highest enrichment (Fig. 1f). This suggests that the positive association between hypermethylation and upregulation in cancer is directly linked to the tissue-specific bivalent chromatin in the pre-cancerous state, rather than being an indirect result of other properties of ESC-bivalent regions. As such, all following analysis was conducted using the tissue-specific histone maps.

Many bivalent genes that become hypermethylated and upregulated in cancer belong to important families of developmental transcription factors, including PAX, SOX, WNT, HOX, DLX, FOX, and ZIC, with known oncogenic roles in cancer (Fig. 1g, Extended Data Figure 1, Supplementary Figure 2). Examples of oncogenic bivalent HyperUp genes include *ZIC5*, which promotes proliferation, aggressiveness, and cancer stemness^34–37^, *SOX2*, a master regulator of cancer stemness^38^, commercially available cancer biomarkers *OTX1*, *ONECUT2* and *TWIST1*^39^, and other genes such as *PAX6*, *WNT2*, *WNT3*, *WNT16*, *DLX5*, and the long non-coding RNA *DLX6-AS1*^40^ (Fig. 1g, Supplementary Table 3). Using CollecTRI^41^, we identified targets of the 84 bivalent HyperUp transcription factors. KEGG pathway analysis on these targets revealed enrichment of many cancer-related pathways, including PI3K-Akt, p53, EGFR, MAPK, FoxO, HIF-1, IL-17, Ras, Wnt, TGF-beta, Hippo, TNF, JAK-STAT, NF-kappa B, Notch, VEGF, and mTOR signaling pathways, as well as pluripotency, apoptosis, cell cycle, senescence and other cancer-relevant pathways (Fig. 1h, Supplementary Table 4). It is therefore important to identify the mechanisms which drive the upregulation of these genes and to understand the role that DNA methylation plays therein.

### Bivalent promoters are enriched in CpGs with positive correlation between methylation and expression

In order to gain a deeper understanding of the link between bivalent chromatin, DNA methylation, and transcription, we computed Spearman correlation between methylation and expression across individuals for each gene in the genome and each CpG site in each of these genes’ promoters (see Methods). Comparing this across cancer types, we observed that a median of 7% of CpG-gene pairs show a significant correlation (defined as q-value < 1e-10, to focus on the strongly correlated pairs, Supplementary Table 1). Of these, the percentage of significant CpG-gene pairs showing negative correlation ranged from 70-94% across the 7 cancer types, with a median of 81%, in line with DNA methylation being primarily a repressive mark (examples shown in Fig. 2a, Supplementary Figure 3). However, the remaining 19% of CpG-gene pairs (4,402 in median) had a positive correlation between expression and methylation, which affected thousands of individual CpGs and over a thousand genes (Supplementary Table 1, examples shown in Fig. 2b, Supplementary Figure 4–5), demonstrating considerable effect of DNA methylation associated with active genes. For example, a CpG probe cg14439629 in the promoter of the *PAX6* gene exhibits a Spearman correlation of R = 0.8 between DNA methylation and *PAX6* expression in the Lung Squamous Cell Carcinoma (LUSC) cancer type (Fig. 2b). Moreover, samples from normal lung tissue have low DNA methylation of this CpG and low *PAX6* expression, compared to high DNA methylation and expression in cancer samples (Fig. 2b), in line with *PAX6* being also a HyperUp gene.

**Fig. 2:**
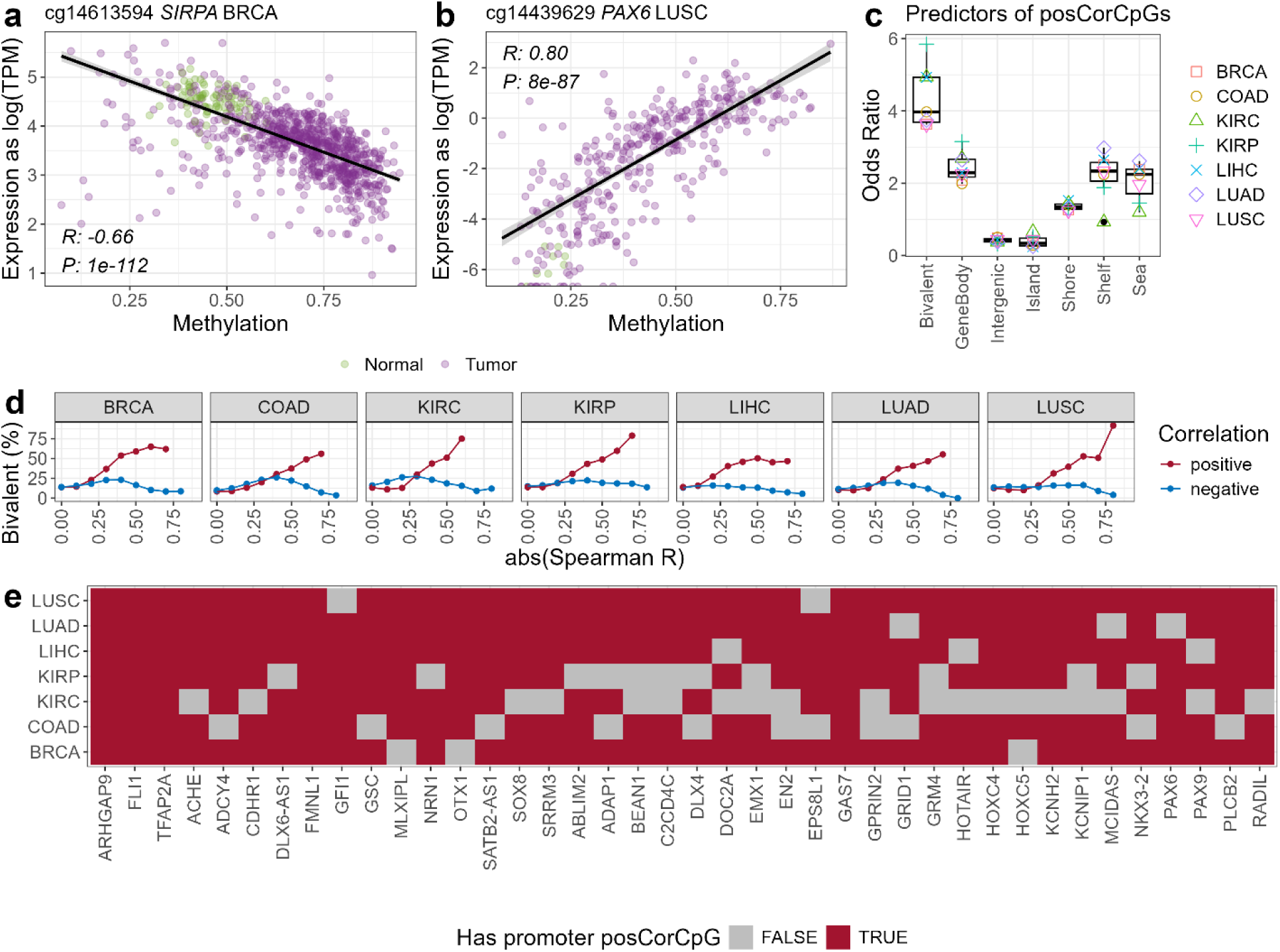
CpG with positive correlation between expression and DNA methylation are enriched in bivalent chromatin. **a,** Example of a negatively correlated CpG-gene pair: a CpG probe in the promoter of SIRPA gene in breast invasive carcinoma (BRCA) patients. Each dot represents one patient sample (normal in green, tumour in purple). Spearman correlation coefficient R and P-value are shown in the bottom left corner. **b,** Example of a positively correlated CpG-gene pair: a CpG probe in the promoter of PAX6 gene in lung squamous cell carcinoma (LUSC) patients. Each dot represents one sample (normal in green, tumour in purple). Spearman correlation coefficient R and P-value are shown in the corner. **c,** Odds ratio of the enrichment of positively correlated CpG-gene pairs in different genomic features (columns), computed in each cancer type (colours). **d,** Percentage of genes with bivalent promoter in matched normal tissue (data from ENCODE) in significantly correlated CpG-gene pairs, stratified by the value of the Spearman correlation coefficient R on the x-axis (positively correlated in red, negatively correlated in blue). **e,** Top 40 genes that are bivalent and have positively correlated CpGs in multiple cancer types.

Next, we compared enrichment of positively correlated CpGs in different genomic features. We observed a significant enrichment of gene bodies in positively correlated CpGs (median odds ratio of 2.3, Fig. 2c), in line with the well-known association between gene bodies and DNA methylation as an active mark^7,15^. CpG islands were overall depleted in positively correlated CpGs, while CpG shelves and open sea were enriched in positively correlated CpGs (Fig. 2c).

The strongest enrichment of positively correlated CpGs was observed for promoters of bivalent genes (Fisher’s exact test OR = 4.0, p < 1e-300; Fig. 2c). This enrichment was reproducible across all tissues, and further increased with higher values of Spearman correlation coefficient R, with as much as 50-80% of highly positively correlated CpGs being in promoters of bivalent genes (Fig. 2d). Moreover, the enrichment of positively correlated CpGs in promoters of bivalent genes was present for CpGs across different categories: inside and outside gene bodies, as well as inside CpG islands, shores, shelves, and open sea (Supplementary Figure 5e). Multiple genes have positively correlated promoter CpGs in multiple cancer types (Fig. 2e).

In summary, our cancer patient data analysis shows that (a) gain of promoter methylation and gene upregulation is a frequent phenomenon across cancer types, (b) the majority of these genes are explained by tissue-specific bivalent chromatin in the matched normal tissue, (c) the mechanisms underlying DNA methylation associated with active transcription is different for gene bodies and bivalent chromatin.

### Bivalent promoters predispose to long-lasting ectopic DNA methylation in epigenome editing

While our observations are very robust, based on thousands of cancer patients and present in all tested cancer types, they represent just a snapshot of the final state, and such data cannot determine causality. We hypothesized that (1) bivalent chromatin is permissive to *de novo* DNA methylation, and (2) this may directly contribute to activation of these originally poised genes. To test these two hypotheses, we turned to epigenome editing.

Epigenome editing allows deposition of *de novo* DNA methylation at CpG sites in a given genomic region by fusing a combination of DNA methyltransferase effectors of DNMT3A and DNMT3L to a DNA targeting module, such as CRISPR/dCas9, engineered zinc finger proteins or TALEs. To ensure long-lasting heritable gain of 5mC, the most successful strategies developed by others and us employ combinations with other chromatin-modifying domains, such as KRAB+DNMT3A+DNMT3L (KAL^42–46^ and CRISPRoff^47^) or EZH2+DNMT3A+DNMT3L (EAL)^44,46^. CRISPRoff contains the same epigenetic domains as KAL but fused to a single dCas9 molecule^47^.

One approach to studying whether hypermethylation of bivalent genes results in gene activation would be to target DNA methylation to specific bivalent genes. However, this approach is low throughput and potentially biased by the choice of the examined genes. That said, in addition to the desired on-target methylation, genome-wide off-target DNA methylation has frequently been reported as a side effect of epigenetic editing with DNMTs alone^48–53^ or in combination with KRAB/EZH2^44^. Here, we leveraged this unintended off-target methylation to identify genomic and chromatin features associated with *de novo* CpG methylation and its effect on gene expression. In particular, this allowed us to assess our two hypotheses in a more unbiased and genome-wide approach.

We therefore investigated global off-target DNA methylation 17 days after hit-and-run epigenome editing, where we transiently expressed the dCas9-based epigenetic editors KAL or EAL targeting the *ERBB2* promoter in the HCT116 colon cancer cell line^44^. *ERBB2*-negative cells were collected by FACS four days after transfection and genome-wide DNA methylation was measured at 17 days after transfection by Illumina Infinium HumanMethylationEPIC (EPIC) array (Fig. 3a). Biological replicates showed strong concordance across the 850,000 CpG probes represented on the EPIC array (R between 0.998 and 0.999, Supplementary Figure 6a-c). For each CpG, methylation levels of KAL or EAL treated cells were compared to control cells treated with dCas9 without effector domain. CpGs with an average methylation increase of at least 0.1 (on the scale 0-1) were considered hypermethylated.

**Fig. 3:**
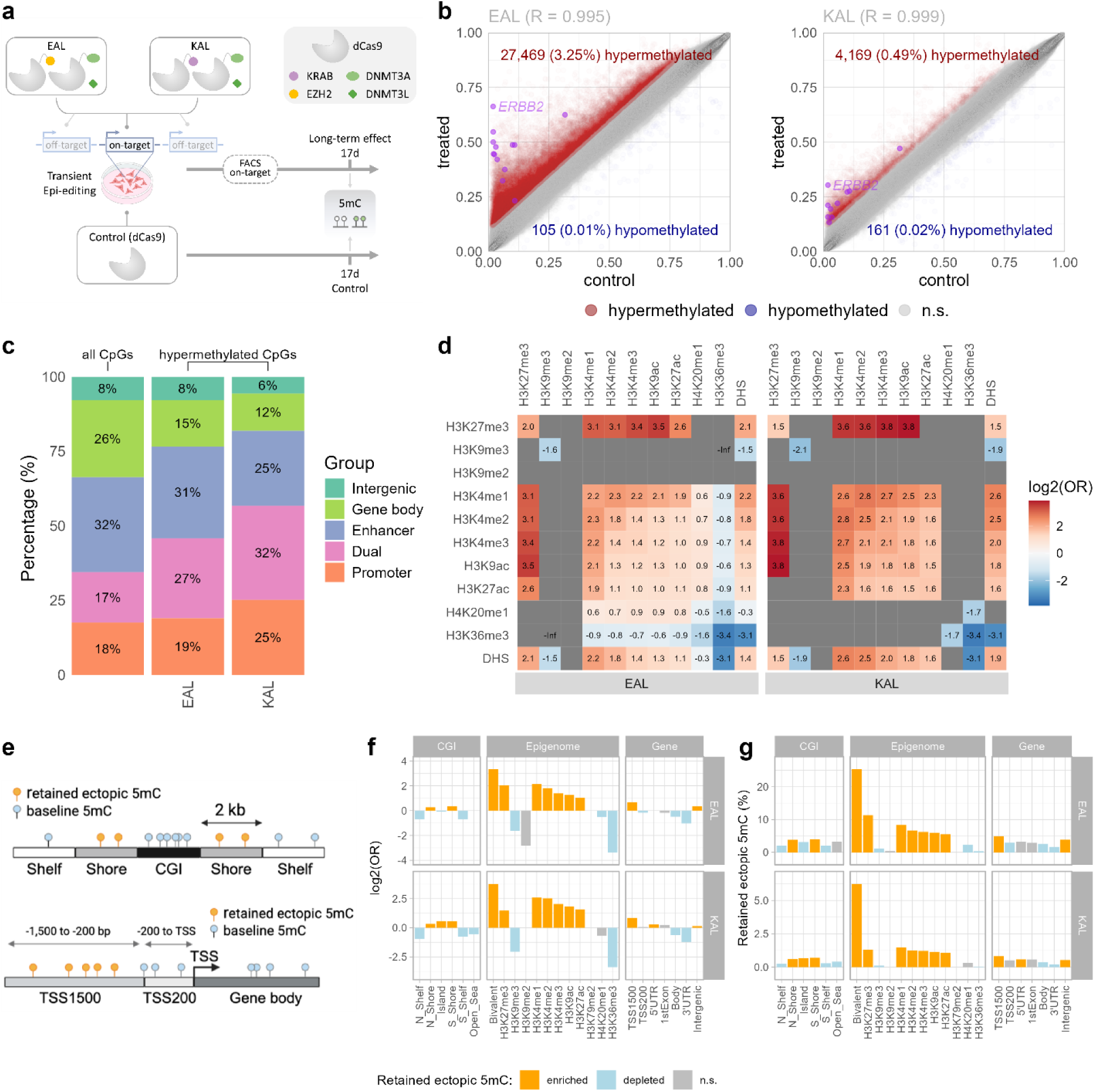
Bivalent chromatin predisposes to long-lasting ectopic DNA methylation in CRISPR/dCas9 epigenome editing. **a,** Diagram of the CRISPR/dCas9 experiment^44^. **b,** Scatterplot showing the CpG methylation values in individual CpGs covered in the EPIC array. Each value represents the average of two replicates. The Pearson correlation coefficient R is shown on top. **c,** Quantification of the EPIC-covered CpGs by the ENCODE-HMM genomic categories^54^. All CpGs represents all CpGs covered in the EPIC array. EAL and KAL represents CpGs with retained methylation 17d after the EAL or KAL treatment, respectively. Dual category represents regions that are both promoters and enhancers (in some cell types). **d,** Log2 odds ratio of the enrichment of retained methylated CpGs in genomic regions marked with a combination of the given two histone marks in HCT116. Significant positive enrichment is shown in red, significant depletion is shown in blue, non-significant combinations are shown in grey. Benjamini-Hochberg (BH) adjusted Fisher’s exact test P-value of at least 0.01 is considered significant. **e,** Diagram of different genomic features used in f-g. **f,** For each genomic category on the x-axis, we show the log2 odds ratio (OR) of enrichment of retained CpG methylation in the given genomic category. Orange represents positive enrichment, blue represents depletion, grey represents non-significant BH-adjusted P-value. **g,** For each genomic category on the x-axis, we show the percentage of CpGs with retained methylation. Orange represents positive enrichment, blue represents depletion, grey represents non-significant BH-adjusted P-value.

It has been suggested that off-target methylation is lost as cells divide. However, we observed that thousands of CpGs retained off-target DNA methylation for at least 17 days (3.2% CpGs for EAL; 0.5% CpGs for KAL, Fig. 3b). Transient delivery ensured that epigenetic editing tools (EAL and KAL) were no longer present 10 days after transfection^43,45^. This demonstrates that some CpGs not only gain methylation, but also retain ectopic methylation even when epigenetic editing tools are no longer present. Over 70% of them fell within CpG probes annotated as ENCODE-HMM promoters or enhancers^54^ (Fig. 3c).

To identify those chromatin marks most enriched in the long-term off-target hypermethylated regions, we used ENCODE measurements of 11 histone marks in HCT116 cells and compared the log2 odds ratio (OR) of hypermethylated probes in all combinations of histone marks (Fig. 3d). Strikingly, the strongest positive enrichment in retained CpG methylation was present in the bivalent combination of H3K27me3 and H3K4me3 (log2 OR 3.4 in EAL and 3.8 in KAL, Fig. 3d-f). The enrichment was similarly high for the combination of H3K27me3 and H3K4me1/2 or H3K9ac, however, the vast majority of these sites also contained H3K4me3, and thus fall within the bivalent category. In total, 25% of bivalent CpGs retained off-target methylation in EAL and 6% in KAL (Fig. 3g). This is more frequent than for any single histone mark alone, or other genomic features (Fig. 3g). This is interesting, because the H3K4me3 mark alone is thought to antagonize DNA methylation^12^. However, our data show that the combination of H3K4me3 and H3K27me3 predisposes for gaining and retaining DNA methylation more than either mark alone.

Next, we verified that the methylation enrichment in bivalent regions was not driven by a confounding correlation with, for example, DNA accessibility, certain genomic features, or initial DNA methylation levels. While retained methylation was in general enriched in CpG island shores (Fig. 3e-g), it was enriched in bivalent regions both inside and outside CpG islands, as well as in CpG island shores and shelves (Supplementary Figure 6d). Similarly, the enrichment of retained methylation in bivalent regions was present in promoters, 5’ UTRs, first exons, gene bodies, as well as intergenic regions (Supplementary Figure 6e). Moreover, it was not explained by the initial DNA accessibility levels (Supplementary Figure 6f) nor baseline DNA methylation levels (Supplementary Figure 6g).

We subsequently repeated the epigenome editing experiment and analysis at a later timepoint, 24 days after treatment, and observed a very similar outcome which further confirmed the durable nature of unintended methylation in bivalent regions (Extended Data Figure 2). Next, we explored whether the off-targets could be explained by dCas9 binding. We observed similar results when using untreated wild type (WT) HCT116 cell line as a control (Extended Data Figure 2) and when using a control of HCT116 cell line treated with dCas9 with no effector domains (Fig. 3). Moreover, we looked at potential dCas9 off-targets based on sequence similarities in the target site. None of the 20 predicted off-targets overlapped with the hypermethylated CpGs (Supplementary Table 5). Together, these observations confirm that the off-targets are not due to dCas9 binding, but rather are a result of presence of DNMTs.

We then wanted to investigate whether retained off-target methylation is specific to our epi-dCas9 epigenome editing approach or whether it is a more general phenomenon. We analysed published whole genome bisulfite sequencing (WGBS) DNA methylation data 7 days after treatment of MCF-7 breast cancer cells with dox-inducible zinc finger protein fused to DNMT3A (Epi-ZF)^55^. In this case, the ZF serves as the DNA binding module, instead of CRISPR/dCas9, and it simultaneously methylates thousands of endogenous promoters. Interestingly, also in the Epi-ZF dataset, retained methylation was enriched in regions originally marked with bivalent chromatin (Supplementary Figure 6h-j).

Finally, we asked whether the identified retained DNA methylation is also enriched in regions which are bivalent in other cell lines, or whether the enrichment is specific to those bivalent regions in the cell line in which the experiment was performed (“matched” cell line). Strikingly, all the EAL, KAL, and Epi-ZF experiments exhibited a clear cell-type specific signal, showing the strongest enrichment of retained methylation in the bivalent regions of the matched cell line (Extended Data Figure 3). This confirms that the association between retained DNA methylation and bivalent chromatin is not driven by other confounding genomic features, such as GC content or the underlying DNA sequence in general.

In conclusion, the epigenome editing data show that bivalent regions predispose for retained DNA methylation by DNMT3A in both Epi-ZF and CRISPR/dCas9 EAL and KAL epigenome editing, and that this is directly linked to the specific properties of the regions marked with the bivalent combination of H3K27me3 and H3K4me3 histone marks.

### CRISPR/dCas9 epigenome editing shows that DNA methylation of bivalent genes leads to their activation

Our above analysis shows that off-target methylation can be widespread but retained CpG methylation is especially enriched in bivalent regions, making this a perfect system to study the effects of DNA methylation gain on gene expression in bivalent vs. non-bivalent chromatin. We performed paired RNA-seq and EPIC array methylation measurements for non-treated HCT116 cells (NT), EAL at 24 days after treatment, and CRISPRoff at 24 days after treatment (Fig. 4). CRISPRoff was included, as it is one of the most popular state-of-the-art epigenome editing tools^47^. To ensure sufficient statistical power to detect also smaller size-effect differences, we performed the RNA-seq measurements in 5 replicates^56^. We detected 9,743 and 432 differentially expressed genes with hypermethylated promoters up to 1,500 bp upstream TSS (“hyper-DE genes”), for EAL (Supplementary Table 6) and CRISPRoff (Supplementary Table 7), respectively. The majority of these genes were downregulated (Fig. 4c,f), in line with promoter methylation being primarily a repressive mark.

**Fig. 4:**
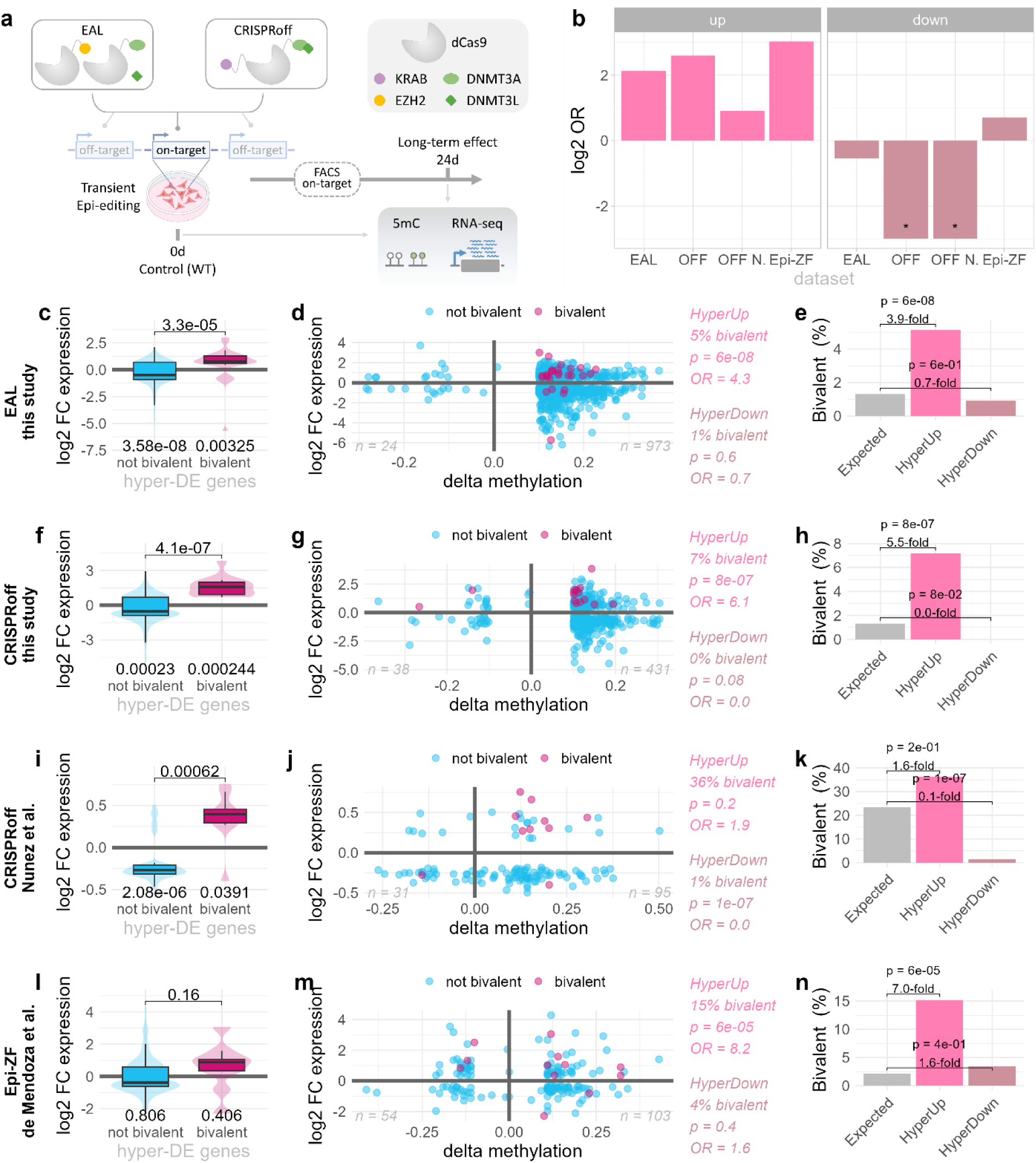
CRISPR/dCas9 epigenome editing shows that hypermethylation of bivalent promoters leads to their activation. **a,** Diagram of the experimental design for the data presented in this figure. RNA-seq and CpG methylation using EPIC array were measured in the non-treated cells and at 24 days after the EAL and CRISPRoff treatments, respectively. **b,** Bivalent genes show enrichment in HyperUp and depletion in HyperDown genes in four datasets: EAL in this study (“EAL”), CRISPRoff in this study (“OFF”), CRISPRoff in the Nunez et al. dataset^47^ (“OFF N.”), and Epi-ZF in the de Mendoza et al. dataset (“Epi-ZF”)^55^. Left: the bars represent log2 odds ratio between HyperUp genes in each dataset, and promoters that are bivalent in the matched cell-line (data on bivalent chromatin from ENCODE). Right: the bars represent log2 odds ratio between HyperDown genes in each dataset, and promoters that are bivalent in the matched cell-line (data on bivalent chromatin from ENCODE). Stars represent values below the y-axis limits (OR 0 for CRISPRoff from this study and OR 1.9 for CRISPRoff from the Nunez et al. dataset^47^). **c,f,i,l,** Violinplot and boxplot visualization of the log2 fold-change (FC) expression in treated/non-treated samples in hyper-DE genes, split by their bivalency status in the given cell line. Non-parametric two-sided Wilcoxon ranksum test P-values are shown (one-sample test in each group compared to zero; and two-sample test to compare the two groups). **d,g,j,m,** Scatterplot showing all differentially methylated and differentially expressed genes in EAL (d) and CRISPRoff (g) in this study, CRISPRoff in the study by Nunez et al.^47^ (j), and Epi-ZF in the study by de Mendoza et al.^55^ (m). The x-axis represents average DNA methylation size effect (treated minus non-treated), while the y-axis represents gene expression size effect (log2 treated/non-treated). Genes bivalent in the matched cell line (data from ENCODE) are highlighted in magenta. In the Nunez et al. dataset, matched cell line data were not available, and therefore bivalency in ESC H1 is shown. The text on the right (top, bright pink, for HyperUp, bottom, dark pink, for HyperDown) represents percentage of genes in that quadrant that are bivalent, as well as Fisher’s exact test P-value and odds ratios of the enrichment of bivalent genes in the given quadrant, compared to expected values based on all genes genome-wide. On-target genes are not shown. **e,h,k,n,** Percentage of bivalent genes: expected (based on all genes) vs. observed in the HyperUp and HyperDown groups. Fold-change of the observed vs. expected values and the corresponding Fisher’s exact test P-value are shown above the bars.

Strikingly, in both EAL and CRISPRoff treated cells, almost all *bivalent* hyper-DE genes were upregulated (Fig. 4c,f, Extended Data Figure 4-5). There are 256 (1.3%) bivalent genes in the HCT116 cell line. By chance, we would therefore expect 1.3% genes in the HyperUp group being bivalent. In EAL, we observed 3.9-fold more than this (*P*-value 6e-8, Fig. 4d-e). In CRISPRoff, this was even greater, with 5.5-fold more bivalent genes than expected (*P*-value 8e-7, Fig. 4g-h). Moreover, the percentage of bivalent genes was non-significantly depleted in the HyperDown group of genes both in EAL (0.7-fold, Fig. 4d-e) and CRISPRoff (0 genes observed, Fig. Fig. 4g-h).

As we observed upregulation upon hypermethylation in bivalent genes also for the CRISPRoff system, we sought to investigate whether a similar effect could be observed with a different target gene and a different cell type. We therefore re-analysed WGBS and RNA-seq data from Nunez et al. where CRISPRoff was used to target the *CLTA* gene in HEK293T cells^47^. The number of differentially expressed genes was lower in the study by Nunez et al., likely because the RNA-seq was performed only in two replicates, limiting the statistical power to detect differentially expressed genes with smaller size effect^56^. Nevertheless, the data confirm this same finding, showing significant upregulation of hypermethylated bivalent genes, and significant downregulation of hypermethylated non-bivalent genes (Fig. 4i-k). Data on bivalency in ESC H1 were used here, as H3K27me3 measurements in HEK293T cells are missing in ENCODE.

Finally, we observed a similar trend also when re-analysing data from the Epi-ZF experiment by de Mendoza et al.^55^, where bivalent genes are also strongly enriched in the hypermethylated upregulated genes (7-fold more than expected, *P*-value 6e-5, Fig. 4l-n).

In summary, we leveraged the unintended off-target methylation of bivalent promoters during epigenetic editing, as analysis of the off-target loci fortuitously provided us with unbiased examples of sufficient statistical power to test (1) whether bivalent chromatin affects gaining or retaining DNA methylation and (2) the resulting effect on gene expression. Our results show that (1) bivalent promoters gain and retain 5mC methylation, and (2) depositing 5mC methylation to promoters of bivalent genes can lead to their activation. The analysis from across the three independent datasets shows that this is reproducible across different epigenome editing tools (EAL, CRISPRoff, Epi-ZF), cell lines (HCT116, HEK293T, MCF-7), target genes (*ERBB2*, *CLTA*, all promoters), duration since treatment (24 days, 28 days, and 7 days), and methylation measurement methods (EPIC array, WGBS).

### Cell-type specificity of bivalent chromatin

Our results show enrichment of bivalent promoters in HyperUp genes and depletion of bivalent promoters in HyperDown genes (Fig. 4). Next, we asked whether this property is specific to the regions that are bivalent in the given cell line as opposed to other cell lines. Strikingly, we observed a clear cell-line specificity in enrichment of bivalent in HyperUp genes (Fig. 5a,c,e). For example, genes bivalent in non-matched cell-lines showed a modest enrichment in genes HyperUp in EAL (odds ratio between 1.2 and 2.6), but the strongest enrichment was present only in genes bivalent in HCT116 (odds ratio 4.3), which is the cell-line where these experiments were performed (“matched cell line”) (Fig. 5a). Similarly, HyperUp genes in CRISPRoff were modestly enriched in non-matched cell lines (odds ratio 0.9-3.3), but very strongly enriched only in the matched cell line (odds ratio 6.1, Fig. 5c). In comparison, the Epi-ZF experiment by de Mendoza et al.^55^ was performed in MCF-7 cell line. In turn, genes that are bivalent in MCF-7 exhibited the greatest HyperUp enrichment (Fig. 5e). Moreover, we also observed strong cell-line specific signal in the depletion of bivalent promoters in HyperDown genes (Fig. 5b,d,f). These results further support that the effect of DNA methylation on preferential gene activation is due to the presence of bivalent chromatin in the given cell line, as opposed to some indirect genomic correlation or other properties of these promoters.

**Fig. 5:**
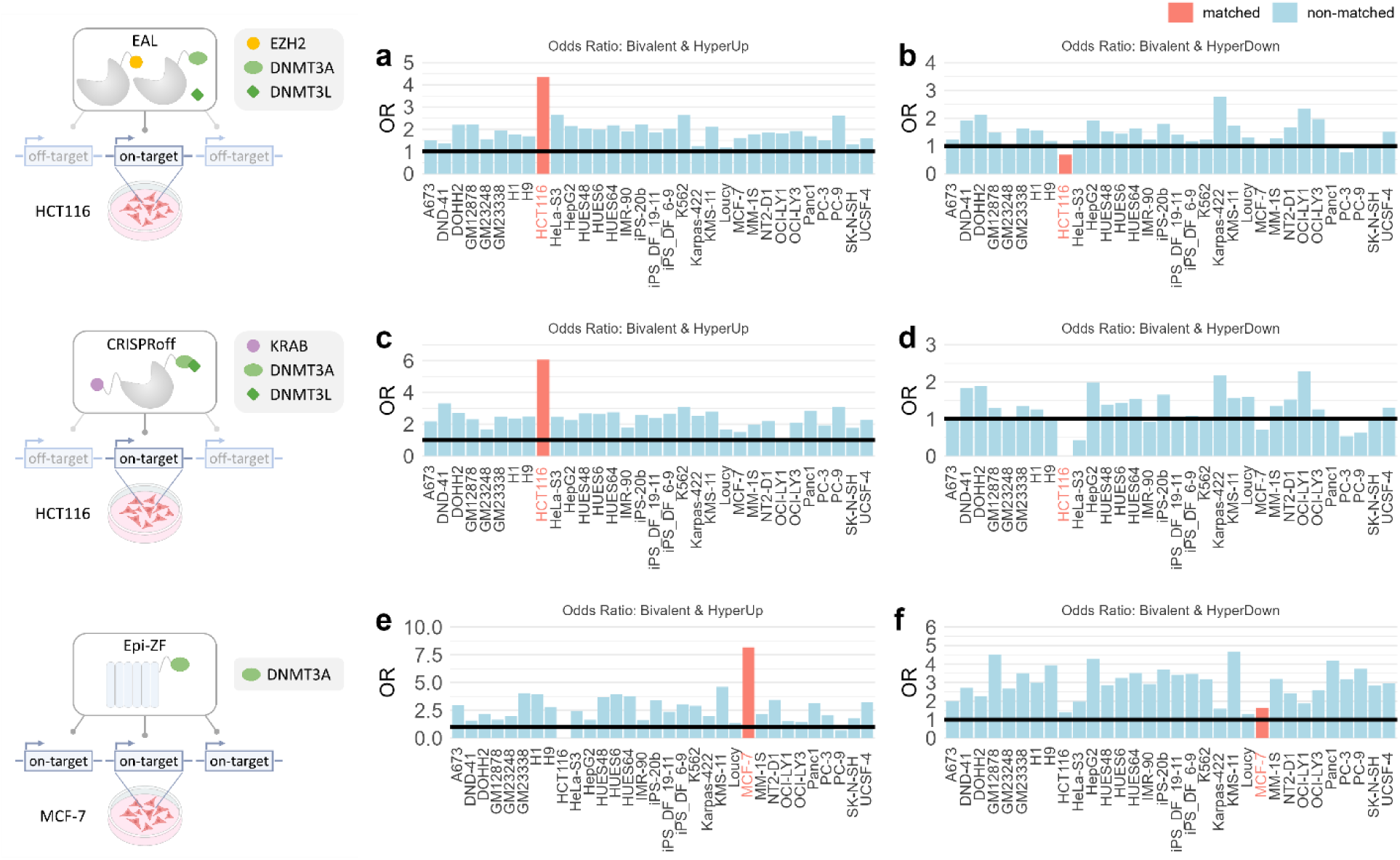
Gene activation upon DNA methylation gain is best explained by cell-type specific bivalent chromatin. a,c,e, Enrichment (odds ratios) between HyperUp (hypermethylated upregulated) genes in EAL (a), CRISPRoff (b), and Epi-ZF de Mendoza et al. dataset^55^, and regions that are bivalent in different cell lines (data from ENCODE). Schematic representations of the experiments are shown to the left of each panel, as relevant. **b,d,f,** Enrichment (odds ratios) between HyperDown (hypermethylated downregulated) genes in EAL (a), CRISPRoff (b), and Epi-ZF de Mendoza et al. dataset, and regions that are bivalent in different cell lines (data from ENCODE). The black horizontal line represents no enrichment (odds ratio 1). The “matched cell line”, i.e., cell-line where the epigenome-editing experiment was performed, is shown in salmon colour, while the other cell lines are shown in blue.

### Loss of MTF2 and H3K27me3 correlate with hypermethylation and activation of bivalent genes

Next, we sought to investigate the mechanism of how DNA methylation may contribute to activate expression of bivalent genes. Bivalent genes are poised because of the repressive mark H3K27me3, which is deposited and maintained by the Polycomb repressive complex 2 (PRC2)^25^. Some of the PRC2 components, most notably MTF2 (also known as PCL2), have been shown to be DNA methylation sensitive and to bind preferentially to unmethylated CpGs^57,58^. Therefore, we hypothesized that 5mC gain in bivalent promoters may block MTF2 binding, and thereby impede maintenance of the repressive bivalent mark H3K27me3, leading to activation (de-repression) of the gene (Fig. 6a).

**Fig. 6:**
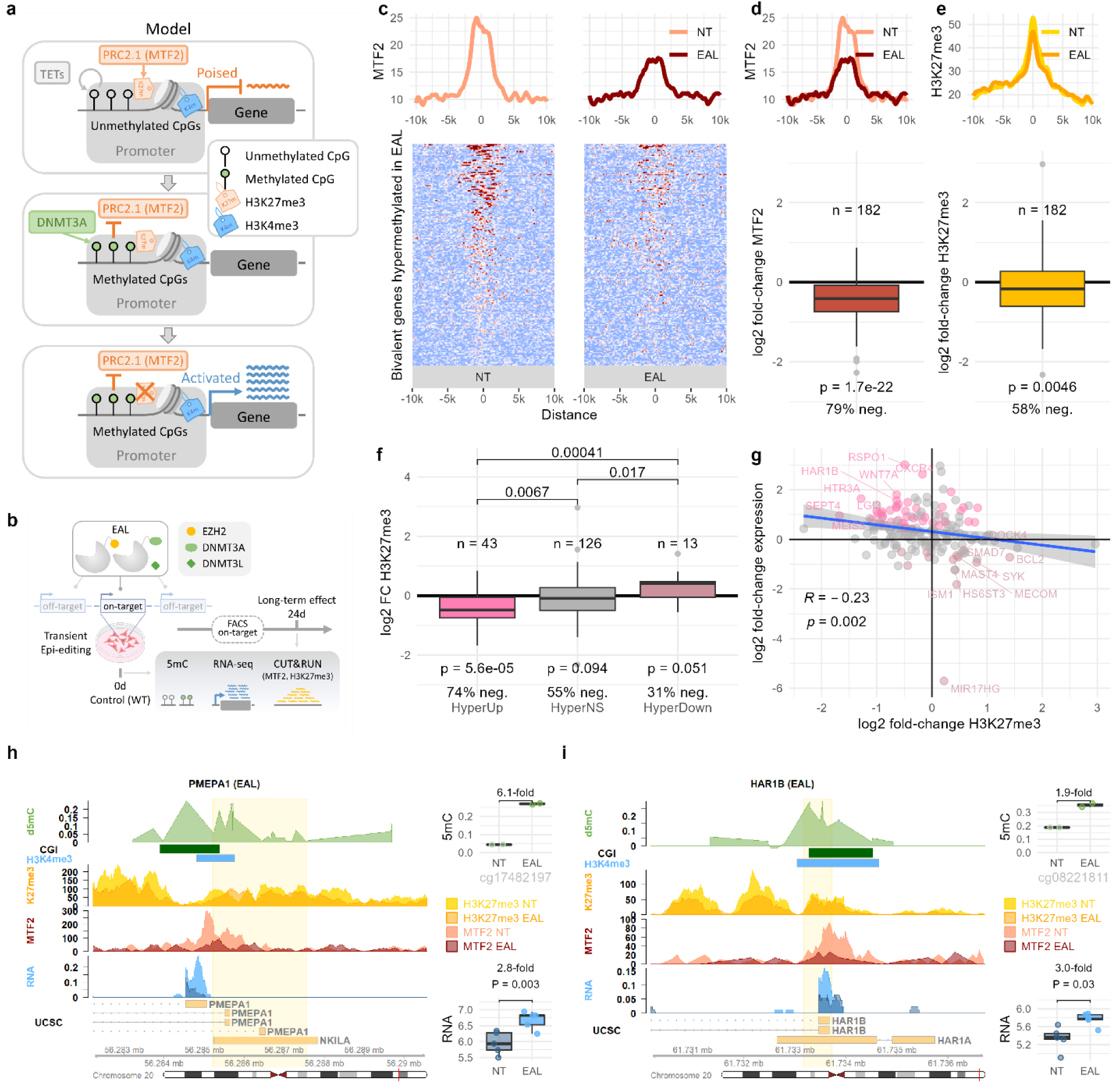
Mechanism: CUT&RUN measurements show reduction of MTF2 and H3K27me3 in hypermethylated upregulated genes 24d after EAL treatment. **a,** Model of the mechanism. Bivalent promoters gain DNA methylation, leading to reduced MTF2 binding, leading to lack of H3K27me3 maintenance (in some genes) and gene activation. **b,** Diagram of the experimental design for the data presented in this figure. MTF2 binding and H3K27me3 occupancy was measured using CUT&RUN in the non-treated cells and at 24 days after the EAL treatment. The data shown in this figure focuses on genes that are bivalent in HCT116 and retained DNA methylation at 24 days after EAL treatment. **c,** MTF2 CUT&RUN signal centered around the bivalent regions in HCT116 of hypermethylated promoters in EAL, shown as a tornado plot heatmap, with the average values shown as a line plot above. **d,** Comparison of MTF2 signal from c in non-treated (NT) and EAL samples shown as a lineplot and a boxplot, representing the distribution of log2(EAL MTF2/NT MTF2) across the 182 bivalent hypermethylated genes. P-value was determined by two-sided paired T-test. **e,** Comparison of H3K27me3 signal from c in NT and EAL samples shown as a lineplot and a boxplot, representing the distribution of log2(EAL H3K27me3/NT H3K27me3) across the 182 bivalent hypermethylated genes. P-value was determined by two-sided paired T-test. **f,** Distribution of log2(EAL H3K27me3/NT H3K27me3) across the 182 bivalent hypermethylated genes, split by their expression status (upregulated in EAL, unchanged, downregulated in EAL). P-values were determined by two-sided paired T-test. **g,** Scatterplot showing log2(EAL H3K27me3/NT H3K27me3) against log2(EAL expression/NT expression) of the 182 bivalent hypermethylated genes. The colours represent differentially upregulated (blue) and downregulated (dark magenta) genes in EAL compared to non-treated samples. The Spearman R and P-value of the correlation are shown in the bottom left corner. **h-i,** Two example bivalent hypermethylated genes in EAL, showing d5mC (EAL – NT), CGI (CpG islands), H3K4me3 peak (in HCT116, from ENCODE), H3K27me3 in NT (light yellow) and EAL (dark yellow) measured by CUT&RUN in this study, MTF2 in NT (light salmon) and EAL (dark red) measured by CUT&RUN in this study, and RNA-seq in NT (dark blue) and EAL (light blue) measured by RNA-seq in this study. The top boxplots show CpG methylation values in the two replicates in the control (NT) and treated (EAL) samples in an example CpG probe (marked with a black dot in the d5mC track). The bottom boxplots show gene-level expression (variance stabilizing transformation values from DESeq2) in the five replicates in the control (NT) and treated (EAL) samples. The DESeq2 adjusted P-value is shown above the boxplot. In the analysis in this figure, hypermethylated genes are defined as genes with at least one CpG with at least 10% higher DNA methylation in EAL compared to non-treated sample (average across two replicates) in the promoter region within distance of up to 1kbp from the promoter bivalent region. In this analysis, upregulated and downregulated genes are defined by adjusted DESeq2 P-value < 0.1.

In order to test this hypothesis, we measured MTF2 binding and H3K27me3 occupancy using CUT&RUN in the non-treated cells vs. EAL and CRISPRoff treatment after 24 days (Fig. 6b, Extended Data Figure 6). In line with our hypothesis, we observed a significant reduction of MTF2 binding in promoters that were bivalent in the non-treated cells and became and remained hypermethylated after treatment (“bivalent hypermethylated genes”) (Fig. 6c-d, Supplementary Table 8-9). This result was reproducible both in EAL (MTF2 reduction in 79% genes, Fig. 6d) and CRISPRoff (MTF2 reduction in 65% genes, Extended Data Figure 6d). The group of all bivalent hypermethylated genes also showed a moderate loss of H3K27me3, but it was not always sufficient to result in gene activation (Fig. 6e).

However, the originally bivalent promoters that became not only hypermethylated but also upregulated (HyperUp), showed a significant reduction in H3K27me3 both after CRISPRoff, as well as EAL treatment (Fig. 6f-g), in line with the loss of H3K27me3 being needed for activation of these genes. This is further supported by the fact that bivalent hypermethylated genes that were not differentially expressed did not show significant change in H3K27me3 levels (Fig. 6f).

To ascertain whether the upregulation is dependent on H3K27me3 reduction, we asked what would happen to bivalent genes that gain 5mC but also gain H3K27me3. We made use of the fact that EZH2, a component of the PRC2 complex important for H3K27me3 deposition, is part of EAL but not of CRISPRoff. Indeed, in EAL but not in CRISPRoff, a minority of bivalent hypermethylated genes became downregulated, and these showed a predominant gain of H3K27me3 (69%, Fig. 6f). This supports the model that the activation of bivalent genes upon 5mC gain is dependent on H3K27me3 reduction.

Finally, we asked whether H3K4me3 is needed for the gene activation upon 5mC gain. We looked at genes that have H3K27me3 but not H3K4me3 in the promoter up to 1500bp upstream or downstream TSS. These “H3K27me3-only” genes were strongly depleted in both HyperUp and HyperDown groups (Extended Data Figure 7). This supports the conclusion that the H3K4me3 mark is needed for activation of bivalent genes upon 5mC gain.

In summary, our data support the following model (Fig. 6a). In non-bivalent genes, gain of DNA methylation in promoters leads predominantly to gene silencing, such as in *SHOX2* and the on-target *ERBB2* (Extended Data Figure 8a-b), in line with the known repressive function of 5mC. However, bivalent regions are susceptible to gain and retain DNA methylation by DNMT3A/L, which leads to reduced MTF2 binding due to its preference for unmethylated CpGs. This often leads to a decrease in H3K27me3, resulting in the activation of originally poised genes, e.g., see *PMEPA1* and *HAR1B* in EAL (Fig. 6h-i) and *RGS9* and *NOG* in CRISPRoff (Extended Data Figure 6h-j). When the loss of H3K27me3 is insufficient, gene expression remains unaffected (Extended Data Figure 8c-d).

## Discussion

Our study has three key findings: (1) bivalent promoters are susceptible to the gain and retention of DNA methylation, (2) gain of DNA methylation in bivalent promoters can lead to their upregulation, and (3) CRISPR/dCas9 epigenome editing demonstrates causality and durability of aberrant *de novo* methylation leading to reduction of H3K27me3 and upregulation of bivalent genes.

As described in many previous studies^21–23^ and observed in our study, bivalent regions are susceptible to hypermethylation during carcinogenesis. This observation has puzzled scientists for decades, as both H3K27me3 and especially H3K4me3 marks were otherwise thought to be mutually exclusive with DNA methylation^12^. In turn, it was proposed that bivalent chromatin protects from DNA methylation^31^ and that it needs to lose H3K4me3^31^ or H3K27me3^59^ before these regions gain methylation. If that was the case, we would expect bivalent regions to be less prone to hypermethylation than regions with just one of the two marks, e.g., H3K27me3. In contrast, we show that bivalent chromatin is directly susceptible to the gain and retention of DNA methylation and that regions marked with both H3K4me3 and H3K27me3 are more prone to hypermethylation than regions with only one of the two marks.

We can speculate on the reasons underlying the susceptibility of bivalent chromatin to gain and retain DNA methylation. The mechanism might involve the previously described recruitment of DNMTs by EZH2^12,60–63^, or another “player” enriched at bivalent regions, such as H2AK119ub, a repressive histone modification deposited by Polycomb repression complex 1 (PRC1). In particular, it was shown that N terminus of DNMT3A1 (the longer isoform of DNMT3A) facilitates enrichment of DNMT3A1 at bivalent promoters through interaction with H2AK119ub^64–68^.

A large body of our knowledge about bivalent regions and about the mutual exclusivity of H3K27me3 and/or H3K4me3 with DNA methylation comes from ESCs^12^. Two mechanisms are likely to contribute to the low DNA methylation levels in bivalent regions in ESCs. First, the DNMT3A isoform expressed in early embryonic development is the short isoform DNMT3A2, which lacks the N terminus domain that facilitates binding to bivalent regions^68^. Second, TET proteins safeguard bivalent promoters from *de novo* DNA methylation in ESCs by oxidising 5mC to 5-hydroxymethylcytosine (5hmC), leading to active DNA demethylation^69^. In turn, loss of this protection, e.g., in TET triple knock outs (TKO) or TET1 deficiency, leads to preferential DNA hypermethylation of bivalent regions^69–71^.

In cancer, active demethylation is also lost, as 5hmC is found at lower levels^72,73^, providing likely explanation why bivalent regions get hypermethylated in cancer^21,29,32,74^. Similarly, when DNMT3A is abundant – as in the epigenome editing experiments – the bivalent regions are readily hypermethylated. A similar scenario occurs when the PWWD domain of DNMT3A is mutated: its binding to H3K36me2/3 is abrogated, and therefore DNMT3A that would normally bind to H3K36me2/3-marked regions such as gene bodies is free to bind to bivalent regions instead, leading to *de novo* methylation in bivalent regions^75–77^.

It has been proposed that replacement of histone modifications by DNA methylation in cancer is to ensure more stable gene repression^7,21,29–32^. However, utilising large scale high-throughput sequencing data, we show that genes that are bivalent in normal tissue and hypermethylated in cancer are in fact predominantly upregulated in cancer. These bivalent genes that we have shown to get hypermethylated and upregulated in cancer include dozens of developmental transcription factors regulating many cancer pathways, such as PI3K-Akt, p53, MAPK, EGFR, apoptosis, senescence, and pluripotency signalling pathways.

We hypothesised that the DNA methylation may directly contribute to the upregulation of these bivalent genes. Our results in several epigenome editing systems demonstrate the causality of aberrant *de novo* methylation at bivalent genes that is sustained and results in gene upregulation. Our CUT&RUN data support a mechanism where the DNA methylation inhibits MTF2 binding of the repressive PRC2 complex, leading to reduction of H3K27me3 and disruption of the repressive chromatin conformation, and resolving the originally poised bivalent promoter into activated state. A recent study showed that targeting histone methyl transferases to deposit H3K4me3 can initiate transcriptional expression on otherwise silent genes^78^. This is in line with our findings that reduction of H3K27me3 at bivalent genes leads to de-repression and gives way to transcriptional activation facilitated by H3K4me3.

Since DNA methylation is typically associated with gene repression, its activating role—albeit within the specific context of bivalent chromatin—may initially seem surprising. However, this is consistent with experimental evidence in example genes and helps explain previous puzzling observations. For example, using Dnmt triple knockout and epigenome editing, Albert et al. elegantly showed that DNA methylation deposition is required for Polycomb eviction and subsequent activation of *Zdbf2*, *Celsr2*, *Atp4a*, and potentially other genes in exit from naïve pluripotency during development^79,80^. In light of our new finding, we investigated the possibility that these genes possess bivalent signatures. We analysed E14 ES cell data from ENCODE and confirmed that all these genes indeed display bivalent signatures, providing further validation of our study. In another example, DNA methylation deposition by DNMT3A is needed for activation of a bivalent gene *PAX6* and for correct morphological and physiological activity in human motor neuron differentiation, as shown using DNMT3A KO and subsequent DNA methylation editing to rescue the *PAX6* DNA methylation and expression and developmental defects^81^. Moreover, in mice carrying a D329A point mutation in the DNMT3A PWWP domain, DNA hypermethylation of bivalently-marked developmental regulatory genes leads to their de-repression^77^. In another example, Dnmt3a-mediated *de novo* DNA methylation at the bivalent *HDAC9* promoter upregulates *HDAC9* expression through repression of H3K27me3 to activate antiviral innate immunity^82^. *De novo* DNA methylation is also needed for activation of the bivalent *FOXA2* transcription factor during endoderm development and differentiation of human ESCs into early endoderm stage cells and pancreatic islet cells^83^. Finally, we have revisited genome-wide studies that reported promoter CpG-gene pairs with positive correlation between DNA methylation and expression, and we showed that the majority of them are explained by bivalent chromatin in the respective tissues (Supplementary Figure 7).

Our results are also in line with studies where TET-mediated promoter demethylation was shown to be required to maintain repressed/poised transcription of bivalent Polycomb-targeted developmental genes in mouse ES cells^70^ and human pluripotent stem-cell-derived human primordial germ-like cells^71^. In particular, deficiency in TET1 was shown to lead to increased DNA methylation in bivalent promoters, decreased PRC2 occupancy, and increased transcription. Similarly, TET TKO in human ES cells exhibit bivalent promoter hypermethylation and within all hypermethylated promoters, the originally bivalent promoters show the highest proportion of upregulated genes^69^.

In conclusion, our study demonstrates that the long-standing view of promoter methylation as purely repressive mark is oversimplified and incomplete. Instead, using epigenome editing, we show that in the context of bivalent chromatin, gain in CpG methylation can instead cause gene activation, with important implications for carcinogenesis and development. We hypothesise that the susceptibility to gain DNA methylation and subsequent activation of the originally poised bivalent genes might be a physiological mechanism used during development, which is hijacked in cancer. As such, it would represent a unique non-mutational epigenetic mechanism driving cancer through activation of bivalent developmental transcription factors that regulate pluripotency, apoptosis, senescence, and other cancer pathways. Given the challenges of direct targeting of transcription factors in cancer, the presented mechanisms might open new avenues of how to target them.

## Methods

### Cancer patient data analysis

TCGAbiolinks (v 2.26.0)^84^ was used to access expression (RNA-seq) and methylation (Illumina Human Methylation 450 BeadChip array) data from The Cancer Genome Atlas (TCGA)^85^. All projects with at least 30 patients with both tumour and normal data and available bivalent map in the matched normal tissue were included (BRCA, COAD, KIRC, KIRP, LIHC, LUAD, LUSC), as summarised in Supplementary Table 1. Tissue-specific and ESC H1 histone mark data for H3K27me3 and H3K4me3 were obtained from the ENCODE^86^ database (Supplementary Table 1). CpG island (CGI) and TSS locations (UCSC genes) with hg19 coordinates were retrieved from UCSC Genome Browser^87^.

For the cancer gene-level differential analysis, default TCGAbiolinks parameters were used for differential expression (edgeR pipeline, glmLRT method, false discovery rate cutoff of 0.01 and log2 fold-change cutoff of 2) and differential methylation (*P*-value cutoff of 1e-5, mean size effect cutoff of 0.25). Promoters were classified as bivalent if they exhibited overlapping H3K27me3 and H3K4me3 marks up to 1,500 bp distant to the TSS. DM-DE genes were defined as differentially expressed genes with at least one differentially methylated CpG in their promoter up to 1,500 bp upstream TSS. Genes with both hypermethylated and hypomethylated CpGs were excluded. For the gene ontology analysis, biological processes and molecular functions were identified using R/Bioconductor clusterProfiler^88^ (enrichGO function) with org.Hs.eg.db^89^, and additional biological pathway analysis was conducted using KEGG pathway analysis^90^ (enrichKEGG from clusterProfiler). Targets of bivalent HyperUp transcription factors were retrieved using CollectTRI^41^. TFBS motifs were identified using ToppFun in ToppGene Suite^91^ (Supplementary Table 3).

For the cancer CpG-level analysis, we extended this cutoff to 3,000 bp both up and downstream the TSS to allow comparison of different genomic features, including gene bodies. Illumina annotation of the HM450K probes (Supplementary Table 10) has been used to classify individual CpG probes by CGI status (Relation_to_UCSC_CpG_Island) and gene body status (UCSC_RefGene_Group). We compiled a list of CpG-gene pairs by taking all genes in the genome and all CpGs that lie in the promoter of that gene. For each CpG-gene pair, we computed the Spearman correlation between DNA methylation and expression across all samples in the given TCGA project. Benjamini Hochberg method was used to correct for multiple testing to define q-values and CpG-gene pairs with q-value < 1e-10 were considered (strongly) significant. Predictors of positively correlated CpG-gene pairs were compared using Fisher’s exact test.

Boxplots throughout the study are computed and plotted R using the function geom_boxplot from the ggplot2 package. The central line represents the median value. The lower and upper hinges correspond to the first and third quartiles (the 25^th^ and 75^th^ percentiles). The upper whisker extends from the hinge to the largest value no further than 1.5 * IQR from the hinge (where IQR is the inter-quartile range, or distance between the first and third quartiles). The lower whisker extends from the hinge to the smallest value at most 1.5 * IQR of the hinge. Data beyond the end of the whiskers are called “outlying” points and are plotted individually.

### Cell culture and transfection

HCT116 (ATCC #CCL-247) were maintained in Dubelco’s McCoy’s 5A media supplemented with 10% FBS and 1% penicillin/streptomycin at 37 °C under 5% CO2. Cells were co-transfected with epi-dCas9 plasmids and a pool of three gRNAs targeting the HER2/ERBB2 gene promoter using Lipofectamine 3000 (Invitrogen) as described previously^44,46^ with a epi-dCas9:gRNA pool amount ratio of 1.25:1. Epi-dCas9 KAL consisted of equal amounts of KRAB-dCas9 (Addgene #112195), DNMT3A-dCas9 (Addgene #100090) and DNMT3L, while EAL included Ezh2[FL]-dCas9 (Addgene #100086), DNMT3A-dCas9 and DNMT3L. Control cells were transfected with the HER2 gRNA pool and dCas9 (no effector domain) (Addgene #100091). Four days after transfection HER2-negative cells were collected by FACS using APC-antihuman CD340 (erbB2/HER-2) (Biolegend #324408), as previously described^44^. Cell sorting was performed using an MoFlo Astrios cell sorter (Beckman Coulter) at the UC Davis Flow Cytometry Shared Resource Core. Each biological replicate was a separate transfection and sorting event. Sorted HER2 negative cells were replated and grown for an additional 20 days (24 days post transfection), harvested at 24 days after transfection and divided for methylation, RNAseq and CUT&RUN analysis.

### DNA methylation

Genomic DNA from two biological replicates per treatment was extracted using Quick-gDNA MiniPrep kit (Zymo Research). The Infinium Human MethylationEPIC BeadChip (Illumina) was used to assess global DNA methylation. DNA samples were processed by the Molecular Genomics Core at the USC Norris Comprehensive Cancer Center. DNA methylation for each CpG was calculated from signal intensities of methylated (M) and unmethylated (U) probes (beta value = M/(M+U)). Background correction and normalization (corrected) was performed with the ‘noob’ function in the minfi software program in R computing language. For each data point, a detection *P*-value was calculated, in which the signal of each analytical probe is compared to a series of background control probes. These *P*-values were incorporated into the beta value files. Beta values that are not significantly different (*P*>0.05) are masked as “NA”. All downstream analysis was conducted using the hg19/GRCh37 human genome assembly. DNA methylation at 17 days after transfection has previously been obtained using EPIC arrays^44^ and was re-analyzed in this study.

Values from the two replicates were averaged and the cutoff of 0.1 between treated and control samples was used to define hypermethylated and hypomethylated CpG probes. All CpG probes were annotated with a range of genomic features: chromatin marks in HCT116 (data from ENCODE), CGI status (Relation_to_UCSC_CpG_Island) and gene body status (UCSC_RefGene_Group) from the Illumina annotation of the HM450K probes, ENCODE-HMM promoters and enhancers^54^, and DNA accessibility levels defines by DNase in HCT116 (Supplementary Table 10). Enrichment of hypermethylated probes in individual genomic features was compared using Fisher’s exact test P-value and odds ratio. To evaluate potential confounding effects, enrichment of bivalent and hypermethylated probes was also computed within restricted sets of probes, such as only in CpG islands, or only in CpG island shores etc (Supplementary Figure 6d-g).

The analysis of Epi-ZF dataset by de Mendoza et al.^55^ (7 days after treatment, MCF7 cell line) was performed in a similar way as above with the following exceptions. The DNA methylation changes were measured using Whole genome bisulfite sequencing (WGBS). Due to the lower precision of WGBS per CpG compared to EPIC arrays^92^, the hypermethylated sites were required to have at least 0.1 difference between the lower of the two treated replicates and the higher of the two control replicates (and vice versa for hypomethylated sites). Moreover, to enable a side by side comparison with our dataset, only CpGs covered by the EPIC array were considered here. Since the experiment was performed in the MCF7 cell line, chromatin marks from the MCF7 cell line (data from ENCODE) were analysed here.

ChopChop (v3)^93^ was used to determine potential SpCas9 off-targets in the human genome allowing up to 3 mismatches for each of the 3 gRNAs targeting the *ERBB2*/HER2 promoter. This identified a total of 20 potential off-target sites (9 for gRNA1, 11 for gRNA2 and 0 for gRNA3).

### RNA-seq

RNA was extracted from five biological replicates using RNeasy Plus Micro kit (QIAGEN). Barcoded strand-specific mRNA libraries were prepared from 400ng total RNA using the NEBNext Ultra II Directional Library Prep kit for Illumina (New England Biolabs). Pooled mRNAseq libraries were sequenced PE150 on the Aviti system (Element Biosciences) at the UC Davis DNA Technologies Core. Sequence reads were then aligned to the human genome (GENCODE GRCh38.p12) using the STAR Universal Aligner^94^ (version 2.7.3a) and the number of reads were counted with the - quantMode GeneCounts option. Differential gene expression analysis was performed with DESeq2^95^ (v 3.6.2) and the adjusted P-value cutoff of 0.01 was used to define differentially expressed genes (DEG), unless stated otherwise.

Promoters were classified as bivalent if they exhibited overlapping H3K27me3 and H3K4me3 marks up to 1,500 bp distant to the TSS. Hyper-DE genes were defined as differentially expressed genes with at least one hypermethylated CpG in their promoter up to 1,500 bp upstream TSS. The average methylation value of hypermethylated or hypomethylated promoter CpGs (up to 1,500 bp upstream TSS) was computed for each gene and plotted in Fig. 4.

### CUT&RUN

For each experiment, 500,000 cells were harvested using TrypLE (Gibco) and cell pellets were washed in Dubelco’s PBS (Gibco). Samples were prepared using the CUTANA ChIC CUT&RUN kit (EpiCyphr) following the manufacturer’s instructions with minor modifications. H3K27me3 was performed with 3-5 replicates, MTF2 was performed with 2 replicates. After immobilizing cells on ConA beads, antibody binding was performed with overnight rotation at 4°C. We used either 1μg H3K27me3 antibody (Millipore 07-0449) or 0.5μg MTF2 antibody (Aviva Systems Biology ARP34292) for final concentrations of 1:50 or 1:100, respectively. 1μg IgG provided in the CUT&RUN kit was used for control assays. HCT116 cells were permeabilized with 0.01% Digitonin and bound chromatin was fragmented and collected using pAG-MNase beads. DNA from antibody bound chromatin fragments was extracted and then purified using Ampure beads (Beckman Coulter). Illumina sequencing libraries were prepared with dual barcodes according to the CUTANA Library Prep Kit (EpiCyphr). Library quality and size distribution was assessed with the LabChip GX (Caliper). Libraries were quantified using the Qubit high sensitivity DNA assay and equal amounts were pooled. Pooled CUT&RUN libraries were sequenced with 75 bp pair-end reads on the Aviti platform (Element Biosciences) yielding 7-9 million reads per library.

H3K27me3 antibody specificity was determined using the SNAP-CUTANA K-MetStat Panel. The fastq files were then filtered using fastp (v 0.23.4)^96^ with default settings. Filtered reads were mapped using bowtie2 (v 2.3.5.1)^97^ with parameters ‘—dovetail --very-sensitive-local -I 10 -X 700’. The duplicates and low-quality reads were removed using MarkDuplicates from gatk^98^ (v 4.1.4.1) and samtools (v 1.9)^99^ with parameters ‘-q 20 -F 1804 -f 2’. Reads overlapping ENCODE DAC Blacklisted Regions ENCFF001TDO were removed using bedtools (v 2.27.0)^100^ intersect. Bam files were converted to bigwig files using deeptools (v 3.5.3)^101^ with parameters ‘--normalizeUsing RPKM --binSize 10 --extendReads --effectiveGenomeSize 2620345972’. Quality control was assessed using deeptools plotFingerprint with parameters ‘--minMappingQuality 30 --skipZeros - -extendReads’. Tornado plot matrices were computed using deeptools computeMatrix with parameters ‘reference-point --referencePoint center -b 10000 -a 10000 --skipZeros’ and plotted using R (v 4.4.2).

All bivalent genes with at least one hypermethylated CpG up to 1,500 bp distant from a TSS (upstream or downstream) have been analyzed in this section. The center of the H3K27me3 peak of the bivalent promoter regions were used for the reference point in the tornado plots. The averaged signal above the tornado plots (including Fig. 6c-e was smoothed using Savitzky-Golay filter (R function sgolayfilt) with parameters p = 3 and n = 151. The gene-level values (e.g., in Fig. 6d-g) were computed as the average signal in the H3K27me3 peak (of the bivalent promoter regions) extended by 1,000 bp on each side. To compare the values between treated and control samples, the log2 fold change was used, i.e., log2(treated/control). In this analysis, upregulated and downregulated genes are defined by DESeq2 adjusted P-value < 0.1, so that more subtle expression changes are also captured (the conclusions were consistent also when the more stringent cutoff of 0.01 was used). The R Gviz package (v 1.42.1)^102^ was used to plot the example genes along genomic coordinates, as in e.g. Fig. 6h-i.

## Acknowledgements

We thank Skirmantas Kriaucionis, Richard White, Yang Shi, and Jakub Tomek for discussions and comments on the manuscript. We thank Alan Jiao for advice and Kate Dunning for language editing. This work was supported by the National Institutes of Health (R21 HG010559-01 to D.J.S. and H.O.), Ludwig Institute for Cancer Research (M.T.), and the Wellcome Trust (225678/Z/22/Z to M.T.). This project was also supported by the UC Davis Flow Cytometry Shared Resources Laboratory with funding from NCI P30 CA093373 and S10 OD018223 and with technical assistance from Bridget McLaughlin, Jonathan Van Dyke and Ashley Karajeh. High throughput sequencing was performed by the DNA Technologies and Expression Analysis Cores at the UC Davis Genome Center, supported by NIH Shared Instrumentation Grant 1S10OD010786-01. EPIC arrays were carried out by the USC Norris Molecular Genomics Core (MGC) supported by Norris Comprehensive Cancer Center CCSG grant (NCI grant # P30CA014089). For the purpose of Open Access, the author has applied a CC BY public copyright licence to any Author Accepted Manuscript (AAM) version arising from this submission.

## Author Contributions Statement

MT and HO jointly conceived the study. HO, MT, and DS designed the experiments. HO performed the experiments with contributions from BB, HY, MM, and CL. AM and MT performed the computational analyses. MT wrote the initial draft, and all authors contributed to the editing of the manuscript. All authors read and approved the final manuscript.

## Data Availability

Sequencing data have been deposited in the Gene Expression Omnibus under accession GSE123830 (EPIC array methylation data from 2019), GSE279655 (EPIC array methylation data), GSE279919 (RNA-seq), and GSE279920 (CUT&RUN data). Other used publicly available resources are listed in Supplementary Table 10.

## Code Availability

All data analysis and code to reproduce figures and tables can be found at https://bitbucket.org/licroxford/cancerbivalentpromoter/ (cancer patient data analysis) and https://bitbucket.org/licroxford/ectopicepiediting (epigenome editing data analysis).

