## Supplementary Materials for "De-novo DNA Methylation of Bivalent Promoters Induces Gene Activation through PRC2 Displacement"

#### Supplementary Tables

Supplementary Table 1: TCGA projects included in the analysis.

Supplementary Table 2: DM-DE genes in the cancer analysis.

Supplementary Table 3: Literature review on bivalent HyperUp genes occurring in at least 3 cancer types.

Supplementary Table 4: KEGG pathway analysis of targets of bivalent HyperUp transcription factors.

Supplementary Table 5: Off-targets as determined by ChopChop (mm3 = max of 3 mismatches).

Supplementary Table 6: Hyper-DE (hypermethylated differentially expressed) genes in EAL.

Supplementary Table 7: Hyper-DE (hypermethylated differentially expressed) genes in CRISPRoff.

Supplementary Table 8: CUT&RUN measurements in bivalent hypermethylated genes in EAL.

Supplementary Table 9: CUT&RUN measurements in bivalent hypermethylated genes in CRISPRoff.

Supplementary Table 10: Used publicly available resources.

### Extended Data Figures

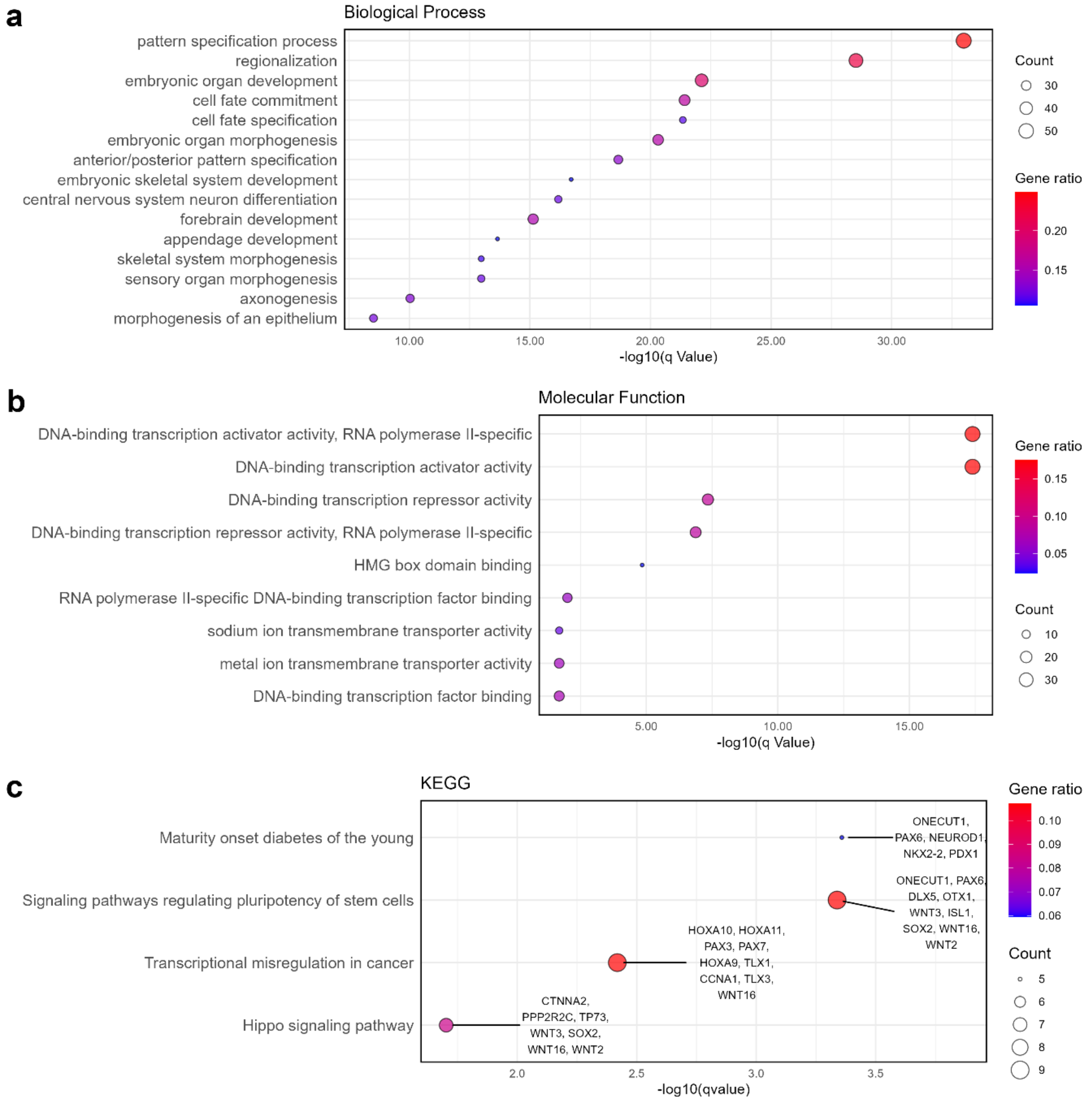

*Extended Data Figure 1: **Gene ontology analysis of bivalent HyperUp genes.** Gene ontology (GO) of Biological Processes **(a)**, Molecular Function **(b)**, and KEGG enrichment analysis **(c)**, was performed using functions enrichGO and enrichKEGG from the Bioconductor package clusterProfiler.*

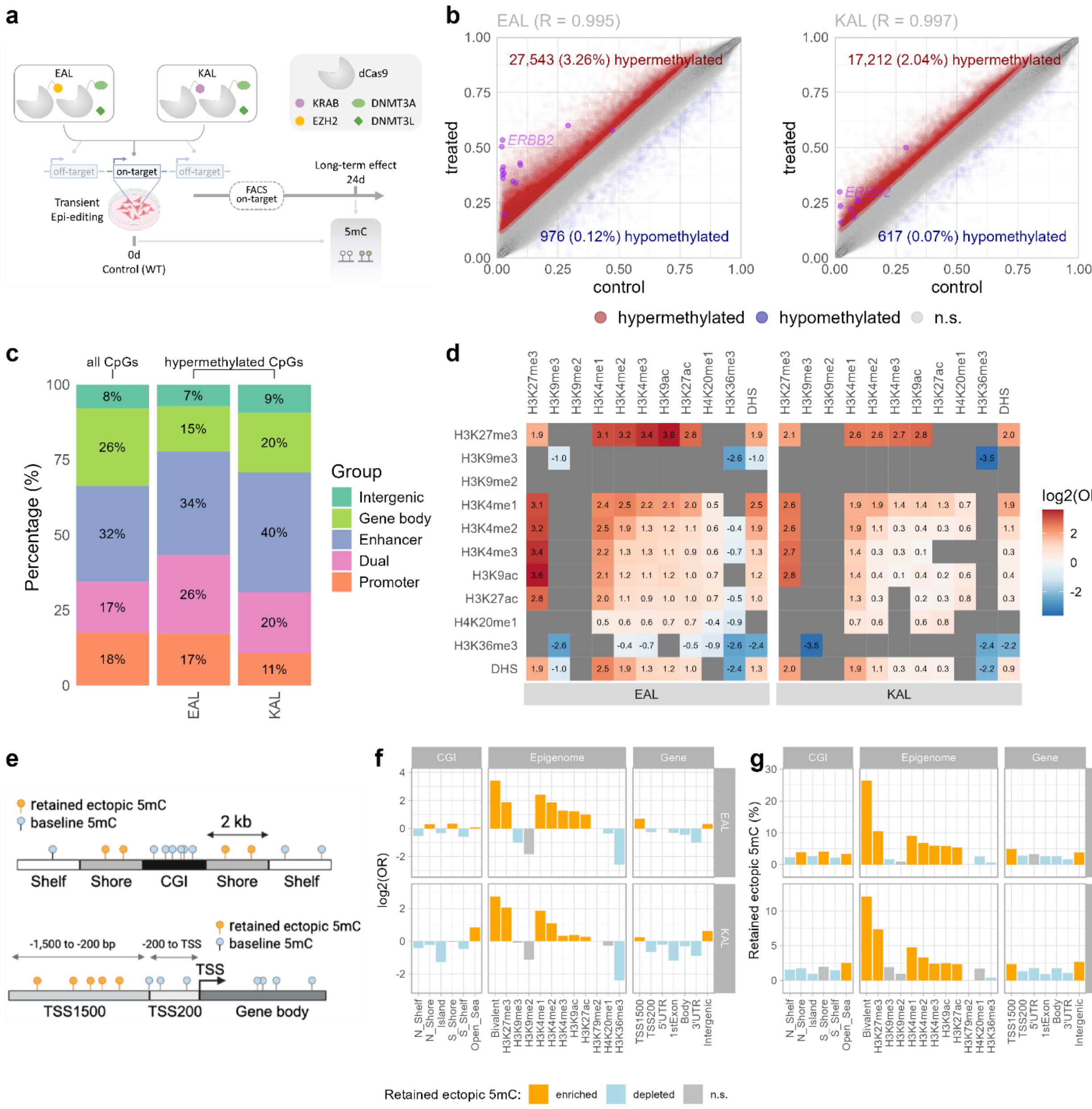

*Extended Data Figure 2:* ***Bivalent chromatin predisposes to long-lasting ectopic DNA methylation in CRISPR/dCas9 epigenome editing.*** *Same as Fig. 3, but a 24-day timepoint, new experiment (from 2024), and a slightly different control (untreated HCT116 cell line as opposed to dCas9-treated 17d timepoint in Fig. 3 control).* ***a,*** *Diagram of the CRISPR/dCas9 experiment.* ***b,*** *Scatterplot showing the CpG methylation values in individual CpGs covered in the EPIC array. Each value represents average of two replicates. The Pearson correlation coefficient R is shown on top.* ***c,*** *Quantification of the EPIC-covered CpGs by the ENCODE-HMM genomic categories. All CpGs represents all CpGs covered in the EPIC array. EAL and KAL represents CpGs with retained methylation 17d after the EAL or KAL treatment, respectively. Dual category represents regions that are both promoters and enhancers (in some cell types).* ***d,*** *Log2 odds ratio of the enrichment of retained methylated CpGs in genomic regions marked with a combination of the given two histone marks in HCT116. Significant positive enrichment is shown in red, significant depletion is shown in blue, non-significant combinations are shown in grey. Benjamini-Hochberg (BH) adjusted Fisher's exact test p-value of at least 0.01*

*is considered significant.* ***e,*** *Diagram of different genomic features used in f-g.* ***f,*** *For each genomic category on the x-axis, we show the log2 odds ratio (OR) of enrichment of retained CpG methylation in the given genomic category. Orange represents positive enrichment, blue represents depletion, grey represents non-significant BH-adjusted p-value.* ***g,*** *For each genomic category on the x-axis, we show the percentage of CpGs with retained methylation. Orange represents positive enrichment, blue represents depletion, grey represents non-significant BH-adjusted p-value.*

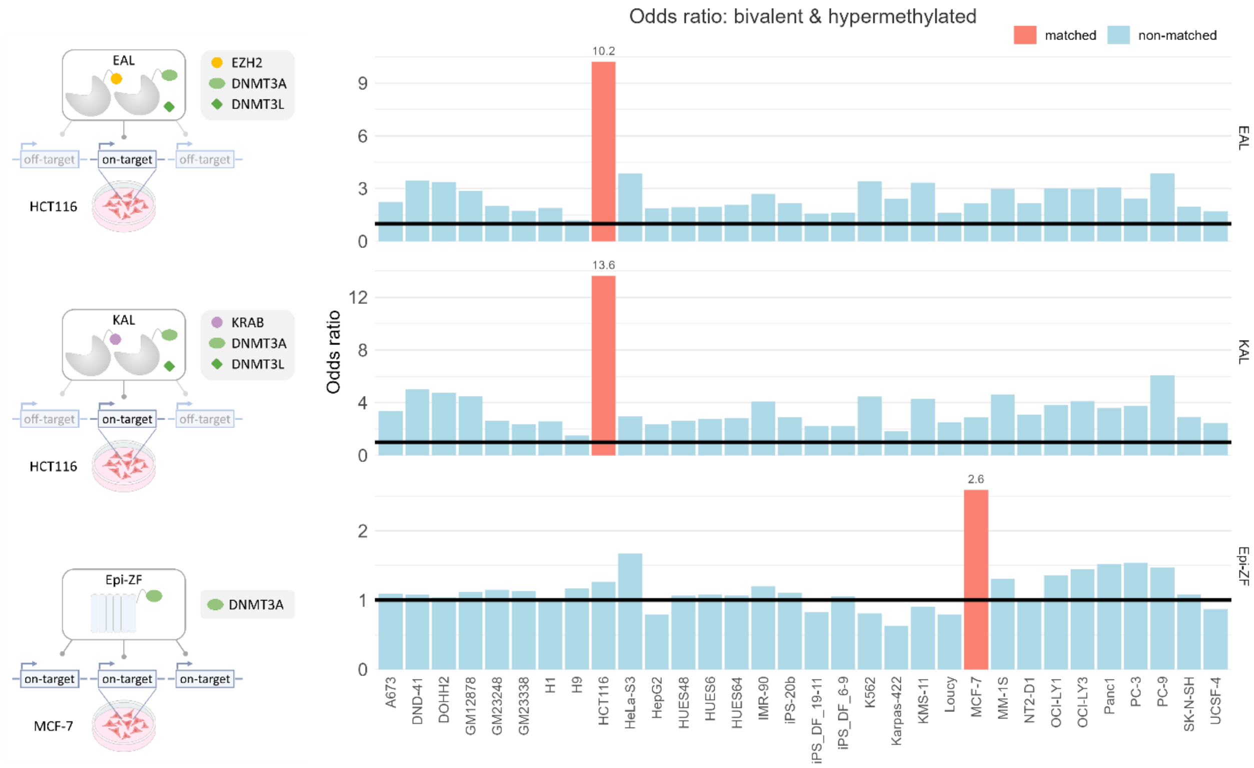

*Extended Data Figure 3:* ***Ectopic methylation is retained in the cell-type specific bivalent regions.*** *Odds ratio (y-axis) of retained CpG methylation in regions marked as bivalent in the given cell line (x-axis). For example, the first bar represents enrichment of retained CpG methylation (in EAL, 17 days post treatment) and regions bivalent in the A673 cell line. The matched cell lines (in salmon colour) represent the cell line, where the experiment was performed. The top two graphs show data from our study (EAL and KAL, both 17 days post treatment), the third graph shows data from the Epi-ZF study by de Mendoza et al. (7 days post treatment withdrawal).*

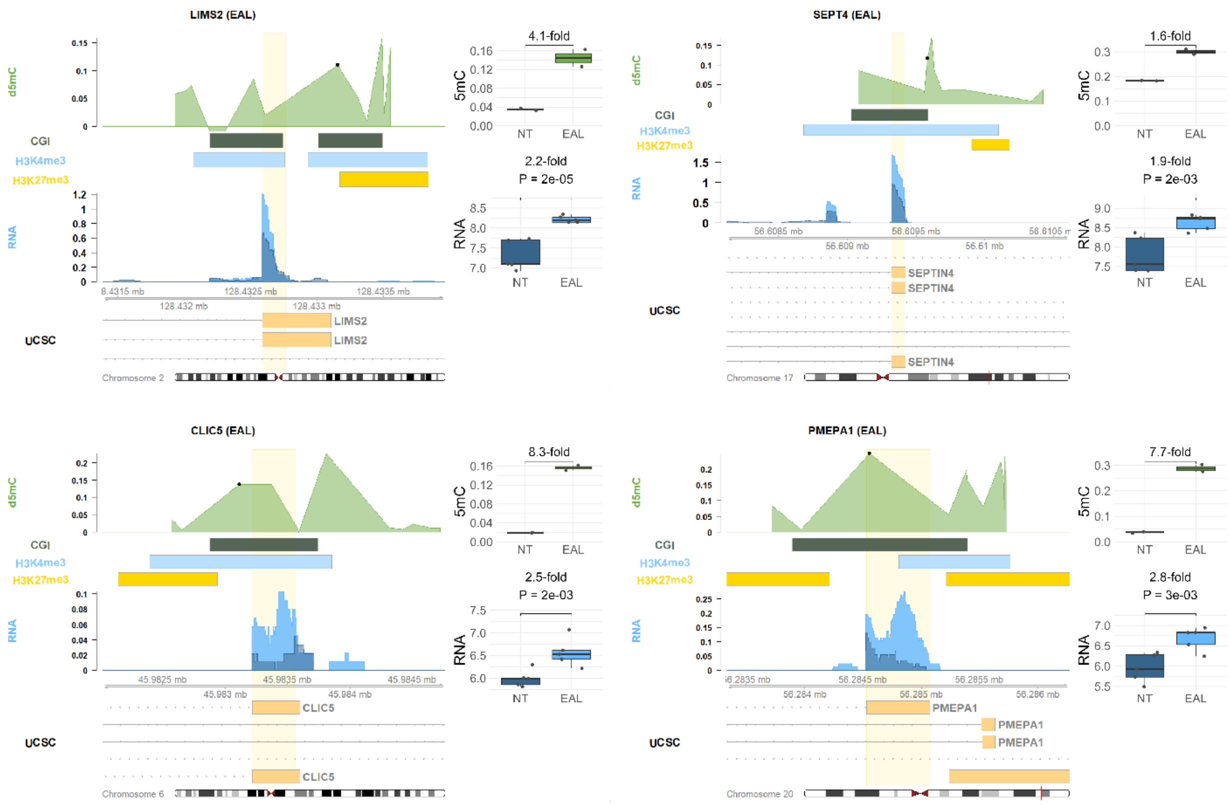

*Extended Data Figure 4:* ***Example genes that are bivalent in HCT116 and HyperUp at 24 days after EAL treatment.*** *d5mC represents the CpG methylation in treated minus control samples (average across two replicates in each). The green boxes represent CpG islands (CGI). The blue boxes represent H3K4me3 peaks in HCT116 (data from ENCODE). The blue boxes represent H3K27me3 peaks in HCT116 (data from ENCODE). The RNA track shows RNA-seq in non-treated (NT, dark blue) and EAL (light blue) measured in this study. The top boxplots show CpG methylation values in the two replicates in the control (NT) and treated (EAL) samples in an example CpG (marked with a black dot in the d5mC track). The bottom boxplots show gene-level expression (variance stabilizing transformation values from DESeq2) in the five replicates in the control (NT) and treated (EAL) samples. The DESeq2 adjusted p-value is shown above the boxplot.*

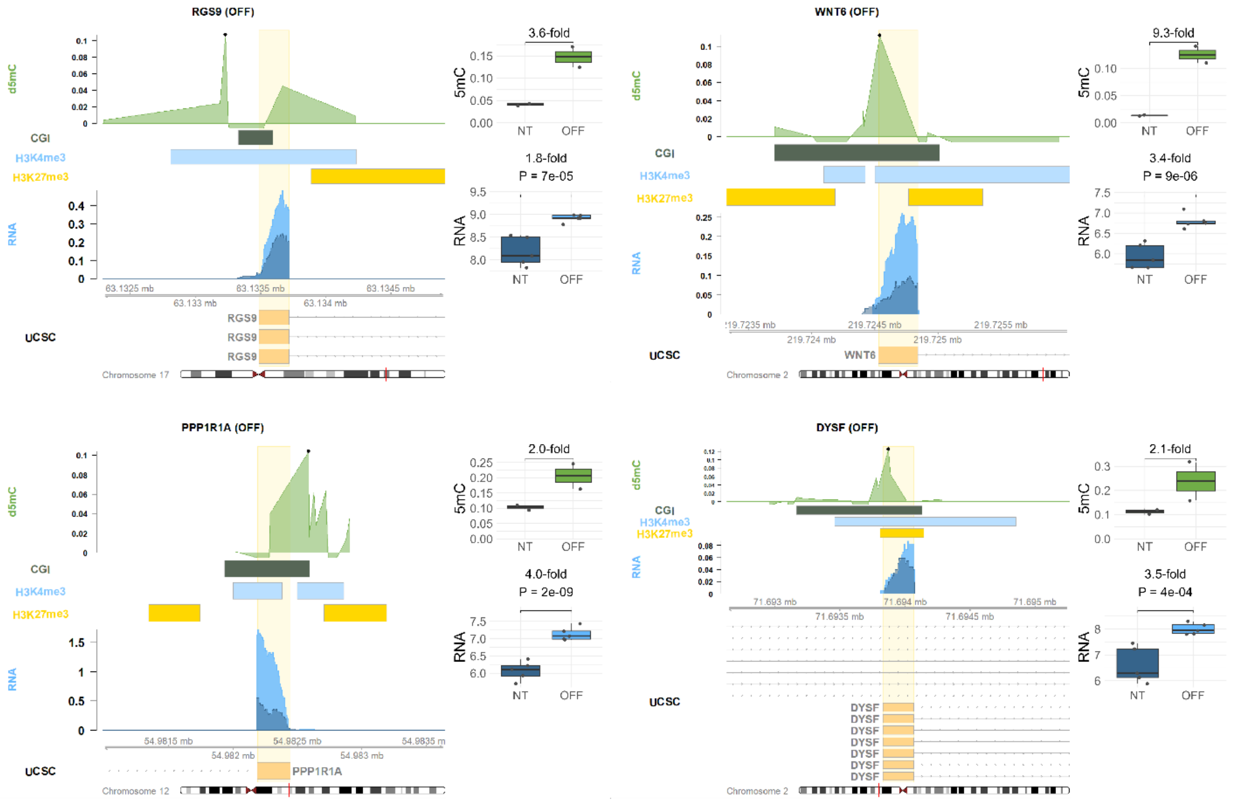

Extended Data Figure 5: ***Example genes that are bivalent in HCT116 and HyperUp at 24 days after CRISPRoff treatment.*** *d5mC represents the CpG methylation in treated minus control samples (average across two replicates in each). The green boxes represent CpG islands (CGI). The blue boxes represent H3K4me3 peaks in HCT116 (data from ENCODE). The blue boxes represent H3K27me3 peaks in HCT116 (data from ENCODE). The RNA track shows RNA-seq in non-treated (NT, dark blue) and CRISPRoff (light blue) measured in this study. The top boxplots show CpG methylation values in the two replicates in the control (NT) and treated (CRISPRoff) samples in an example CpG (marked with a black dot in the d5mC track). The bottom boxplots show gene-level expression (variance stabilizing transformation values from DESeq2) in the five replicates in the control (NT) and treated (CRISPRoff) samples. The DESeq2 adjusted p-value is shown above the boxplot.*

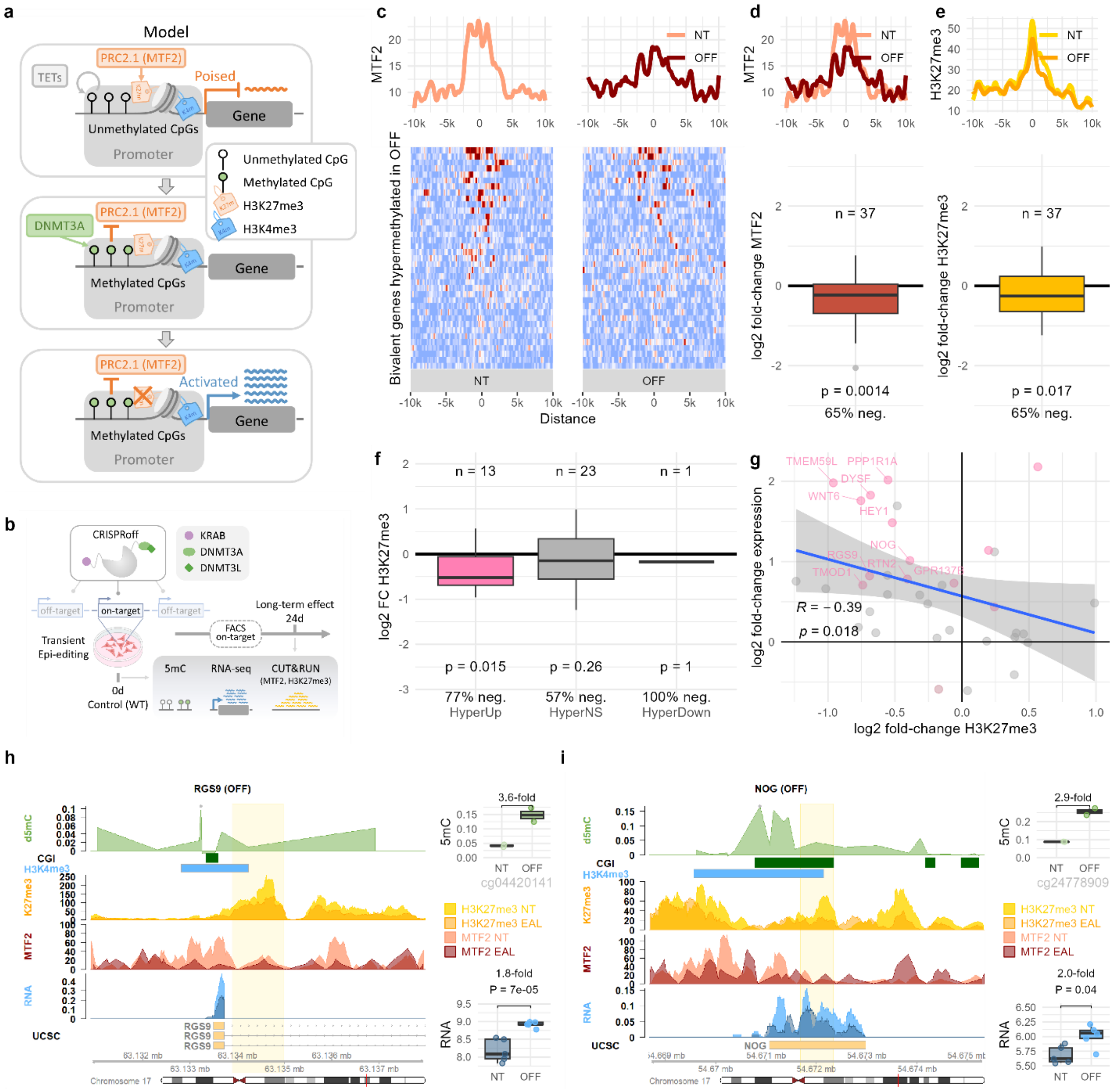

*Extended Data Figure 6:* ***Mechanism: CUT&RUN measurements show reduction of MTF2 and H3K27me3 in hypermethylated upregulated genes 24d after CRISPRoff treatment.*** *The same as Fig. 6 but for CRISPRoff (shortened as "OFF").* ***a,*** *Diagram of the experimental design for the data presented in this figure. MTF2 binding and H3K27me3 occupancy was measured using CUT&RUN in the non-treated cells and at 24 days after the EAL treatment. The data shown in this figure focuses on genes that are bivalent in HCT116 and retained DNA methylation at 24 days after EAL treatment.* ***b,*** *Model of the mechanism. Bivalent promoters gain DNA methylation, leading to reduced MTF2 binding, leading to lack of H3K27me3 maintenance (in some genes) and gene activation.* ***c,*** *MTF2 CUT&RUN signal centered around the bivalent regions in HCT116 of hypermethylated promoters in EAL, shown as a tornado plot heatmap, with the average values shown as a line plot above.* ***d,*** *Comparison of MTF2 signal from c in non-treated (NT) and EAL samples shown as a lineplot and a boxplot, representing the distribution of log2(EAL MTF2/NT MTF2) across the 182 bivalent hypermethylated genes. P-value was determined by two-sided paired T-test.* ***e,*** *Comparison of H3K27me3 signal from c in NT and EAL*

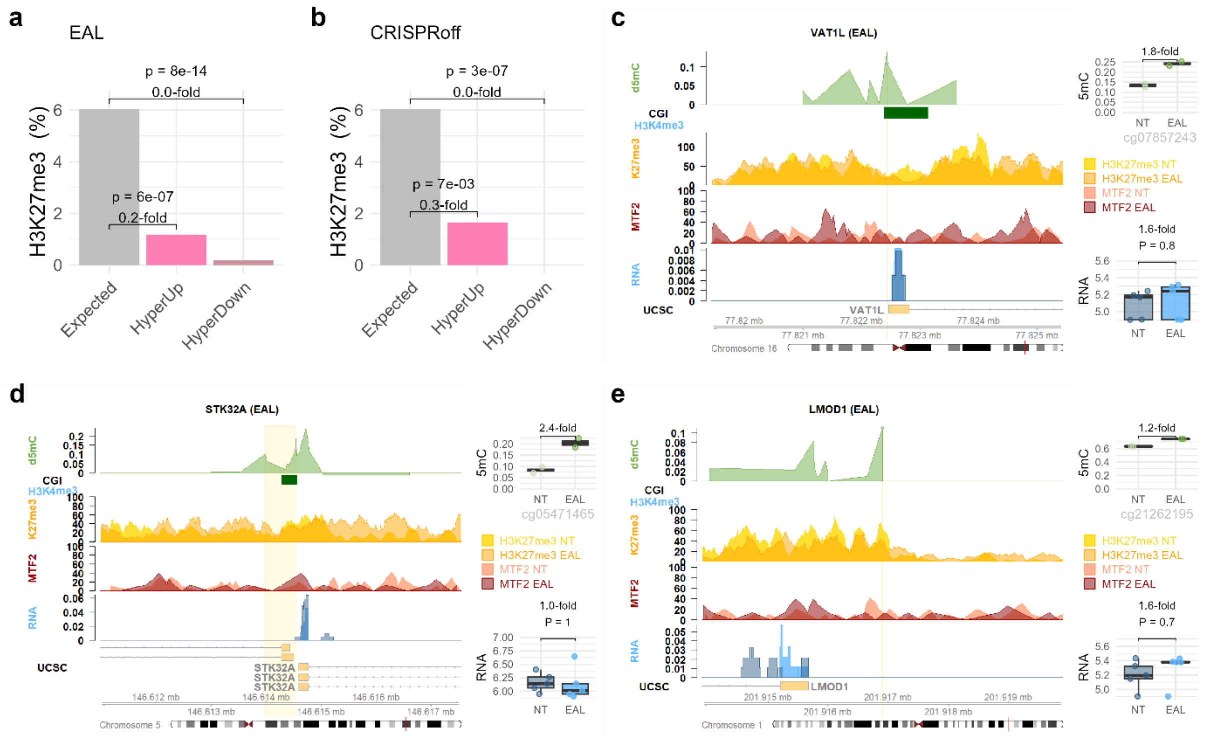

*Extended Data Figure 7:* ***Promoters marked with H3K27me3 but not H3K4me3 do not get activated upon 5mC gain. a-b,*** *H3K27me3-only genes are depleted in HyperUp and HyperDown groups both in EAL (a) and CRISPRoff*

*(b) treatments. The bars show percentage of H3K27me3-only genes (i.e., lacking H3K4me3): expected (based on all genes) vs. observed in the HyperUp and HyperDown groups. Fold-change of the observed vs. expected values and the corresponding Fisher's exact test P-value are shown above the bars.* ***c-e,*** *Example genes that gain 5mC, and are marked with H3K27me3, but not H3K4me3.*

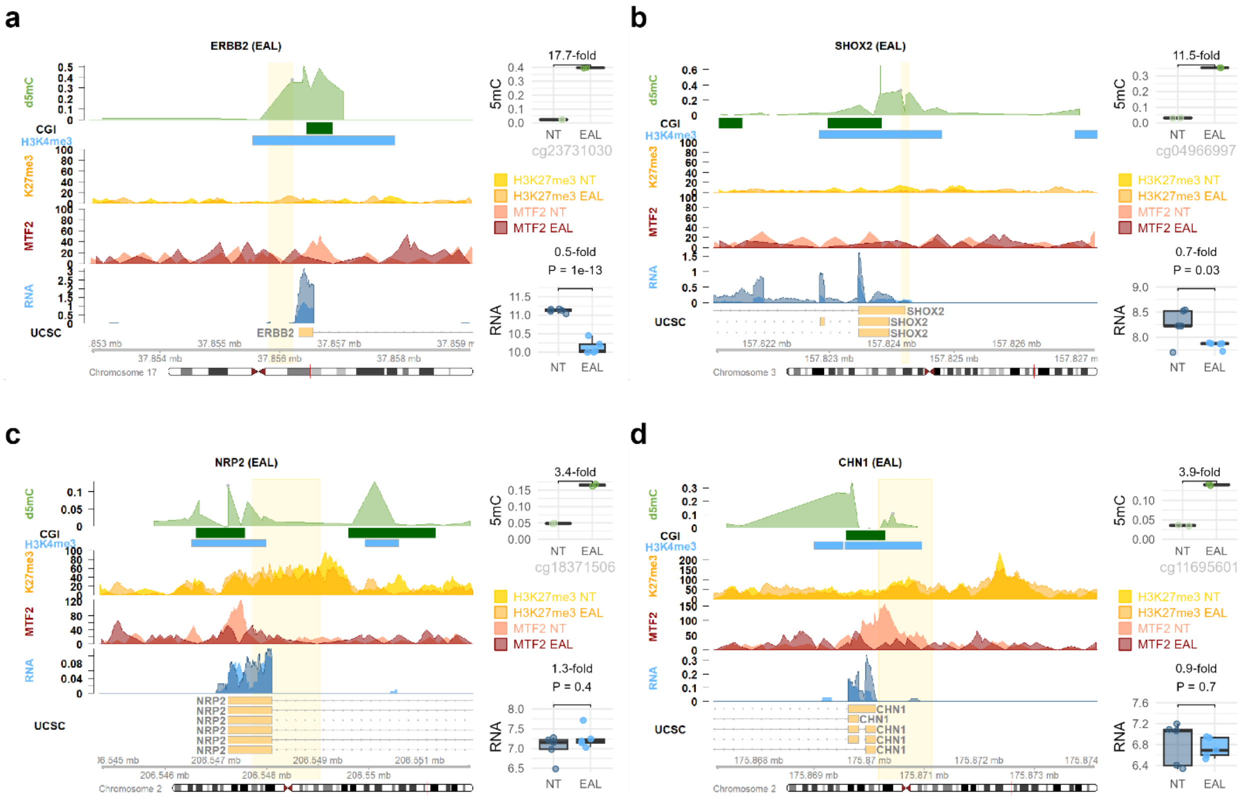

*Extended Data Figure 8:* ***CUT&RUN data for EAL in example genes: on-target non-bivalent gene ERBB2 (a), off-target non-bivalent gene SHOX2 (b), bivalent genes with insufficient H3K27me3 decrease and lack of differential expression (c-d).*** *The tracks in the genomic visualisation are: d5mC (EAL – NT), CGI (CpG islands), H3K4me3 peak (in HCT116, from ENCODE), H3K27me3 in non-treated (NT, light yellow) and EAL (dark yellow) measured by CUT&RUN in this study, MTF2 in NT (light salmon) and EAL (dark red) measured by CUT&RUN in this study, and RNA-seq in NT (dark blue) and EAL (light blue) measured by RNA-seq in this study. The top boxplots show CpG methylation values in the two replicates in the control (NT) and treated (EAL) samples in an example CpG probe (marked with a black dot in the d5mC track). The bottom boxplots show gene-level expression (variance stabilizing transformation values from DESeq2) in the five replicates in the control (NT) and treated (EAL) samples. The DESeq2 adjusted p-value is shown above the boxplot.*

### Supplementary Figures

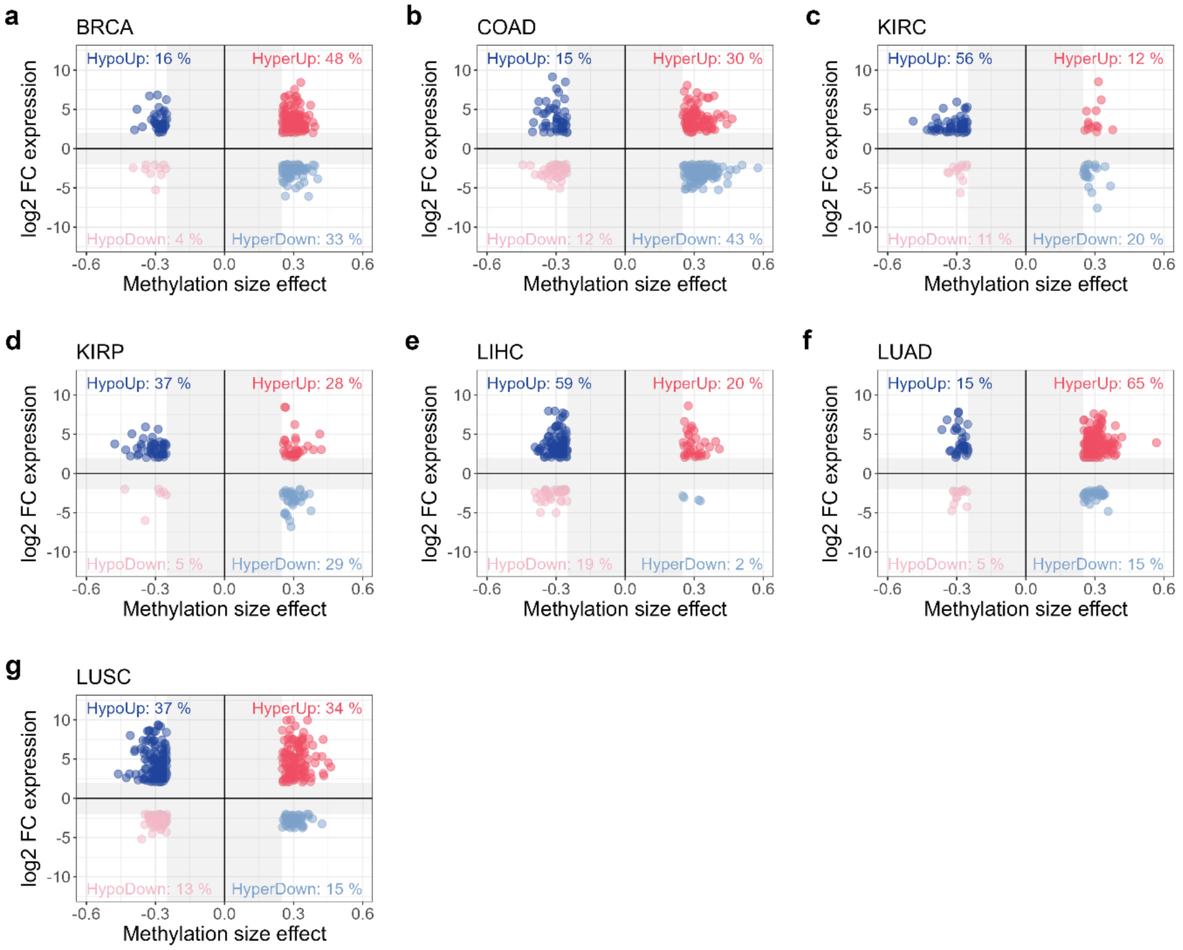

*Supplementary Figure 1:* ***Hypermethylated and upregulated genes are frequent in cancer and largely explained by bivalent chromatin in matched normal tissue, consistently across all studied cancer types*****. a-g,** *All genes that are differentially methylated and differentially expressed in the given cancer type. The x-axis shows methylation size effect (hypermethylated in cancer on the right), computed as average cancer minus normal CpG methylation of the differentially methylated promoter CpGs of that gene. The y-axis shows expression size effect (upregulated in cancer on the top), computed as $log_2$ fold-change of expression in cancer/normal. Promoter is defined as up to 1500 bp upstream TSS.*

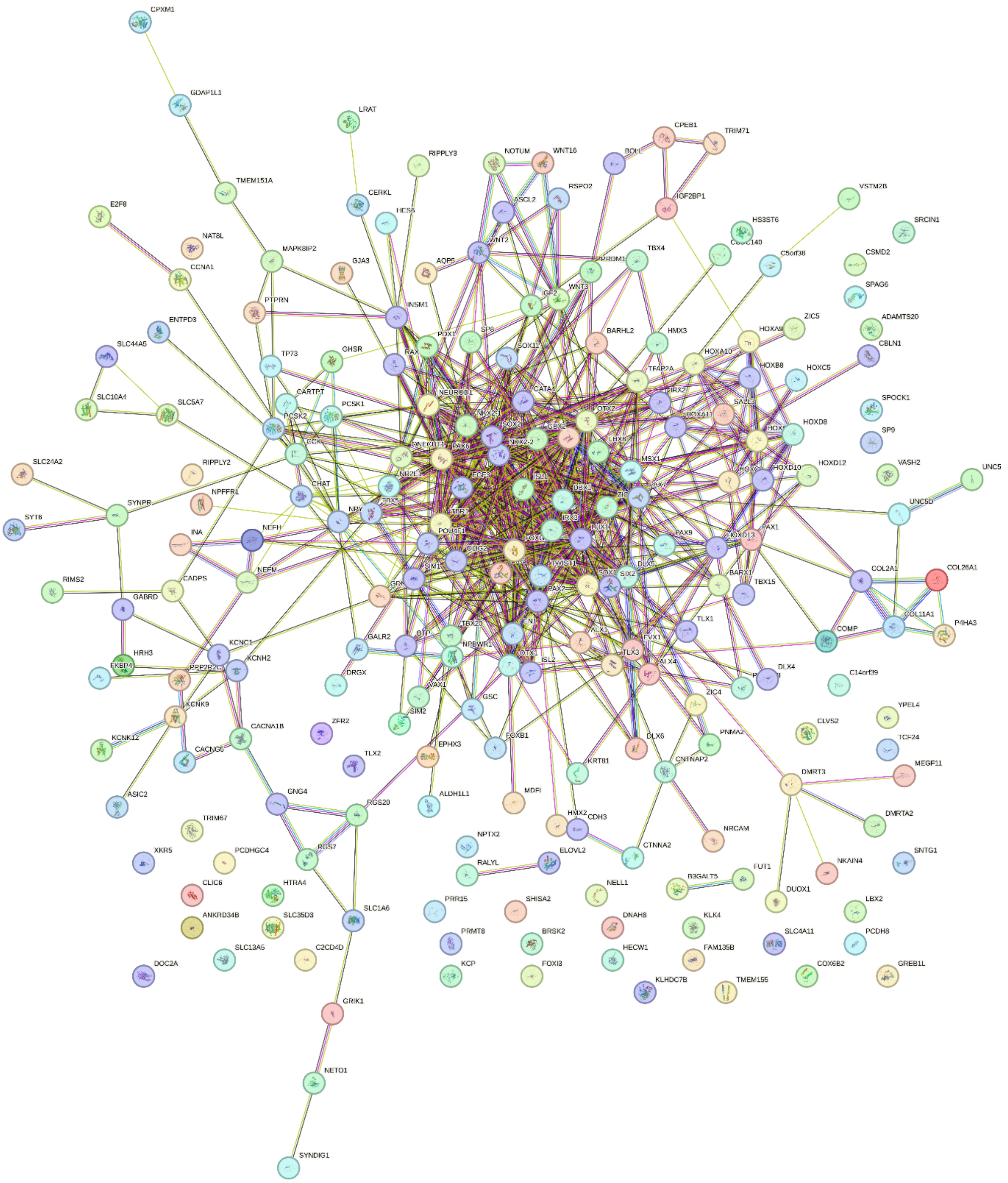

*Supplementary Figure 2:* ***Gene interaction network of bivalent HyperUp genes generated using STRING.*** *Network nodes represent proteins. Empty nodes represent proteins of unknown 3D structure, filled nodes have a known or predicted 3D structure. Edges represent protein-protein associations: known interactions from curated databases (light blue) or experimentally determined (purple); predicted interactions by gene neighbourhood (dark green), gene fusions (red), or gene co-occurrence (dark blue); and others including text-mining (light green), co-expression (black), or protein homology (lilac). Default analysis settings were used. Of the 233 bivalent HyperUp genes, 221 that were present in the database are shown.*

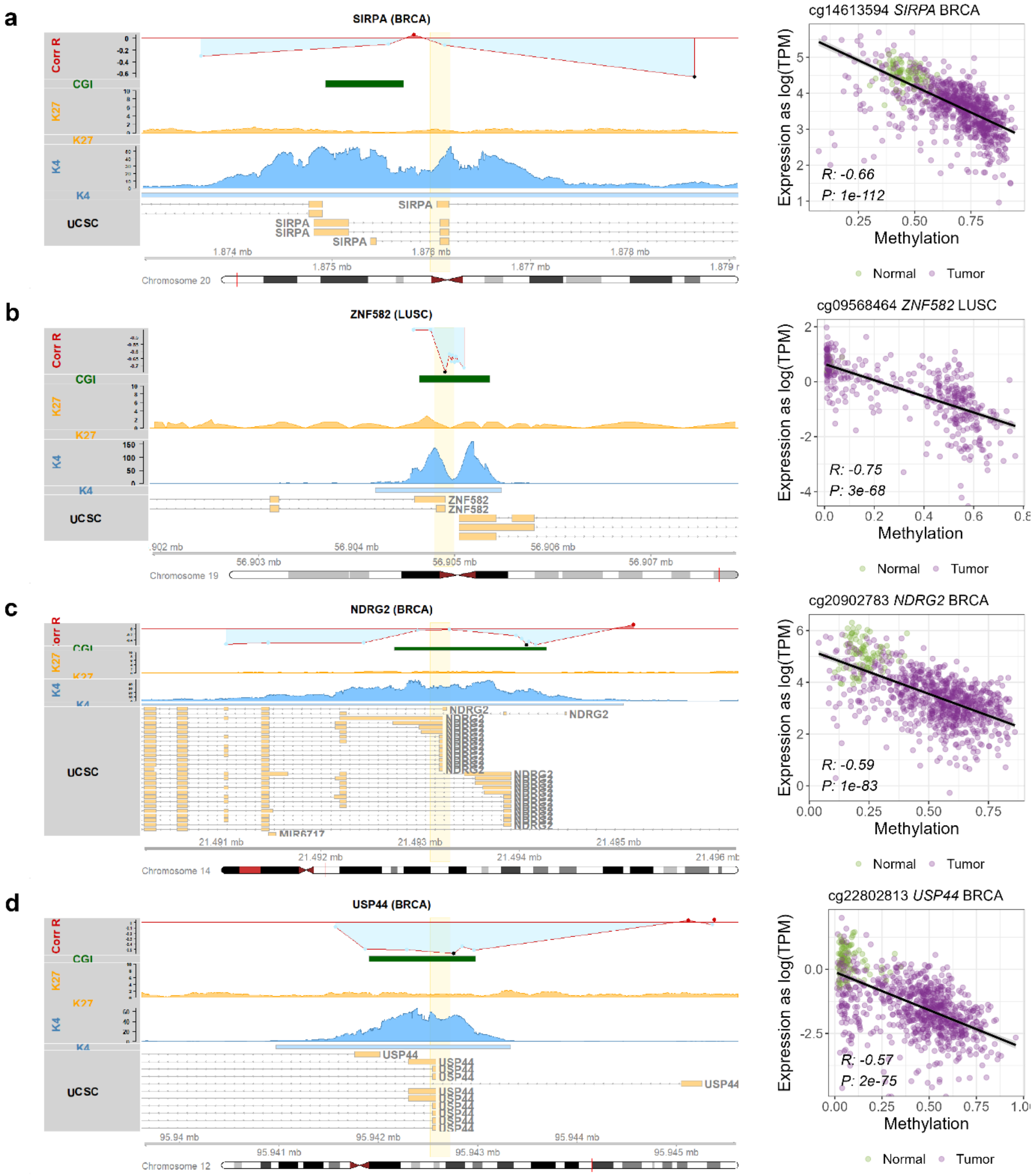

*Supplementary Figure 3:* ***Examples of genes with strongly negatively correlated CpGs in their promoter regions. a-d,*** *Left: Genomic visualization (hg19) of different (epi)genomic features in the example promoter regions: Corr R represents the Spearman Correlation coefficient between DNA methylation in the given CpG and gene expression of the given gene, with positive correlation shown in red and negative correlation shown in light blue. The black dot represents the CpG shown in the scatter plot on the right. CGI represents a CpG island. K27 represents H3K27me3 in the matched normal tissue, shown as signal track (filled polygon) shown above called peaks (box). K4 represents H3K4me3 in the matched normal tissue, shown as signal track (filled polygon) shown above called peaks (box). UCSC track represents gene transcripts obtained from UCSC. Right: Scatterplot of the DNA methylation (x-axis) and gene expression (y-axis) for the given CpG probe and the given gene. Each*

*dot represents one sample (normal in green, tumour in purple). Spearman correlation coefficient R and p-value P are shown in the bottom left corner.*

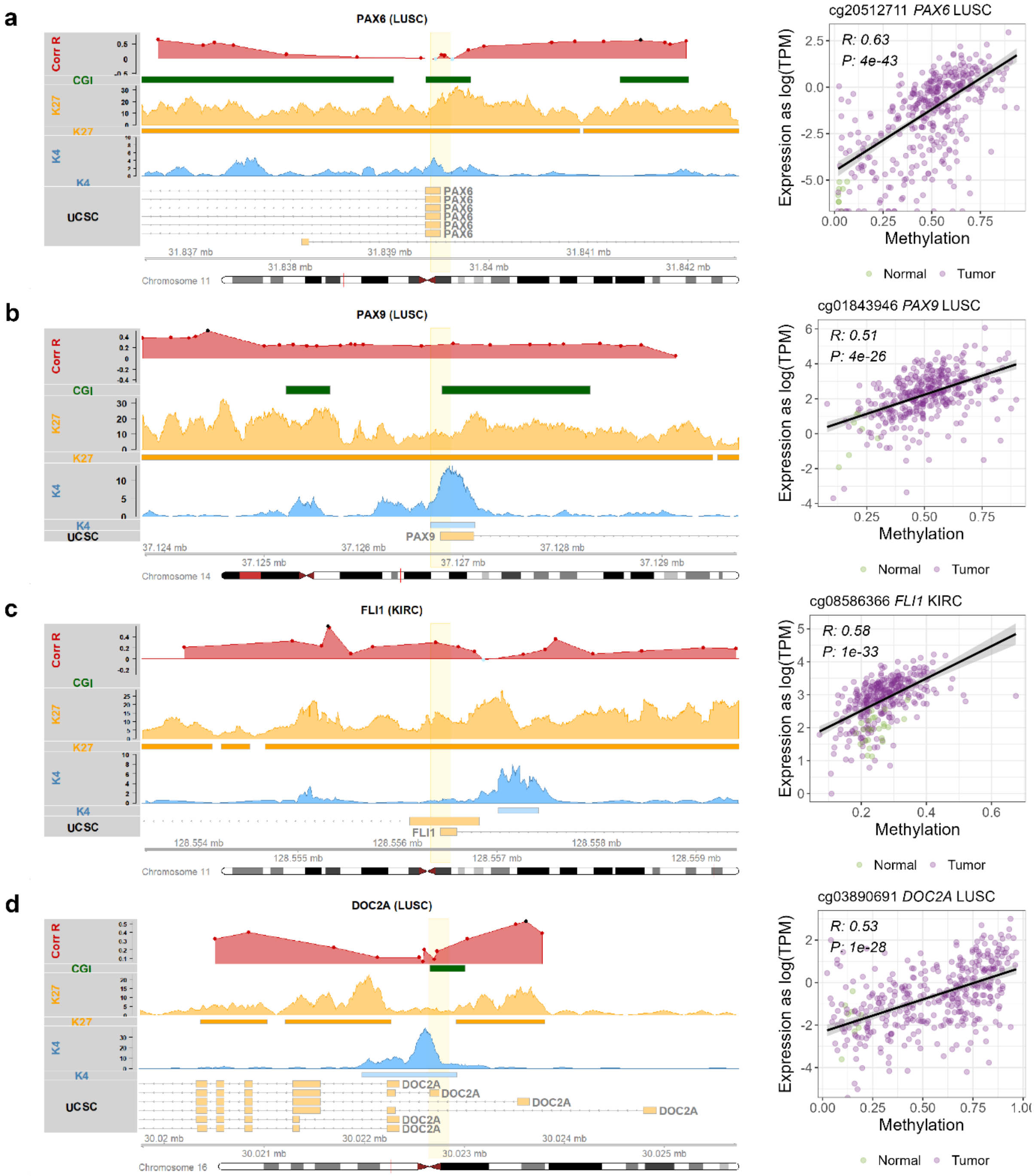

*Supplementary Figure 4: **Examples of bivalent genes with strongly positively correlated CpGs in their promoter regions. a-d,** Left: Genomic visualization (hg19) of different (epi)genomic features in the example promoter regions: Corr R represents the Spearman Correlation coefficient between DNA methylation in the given CpG and gene expression of the given gene, with positive correlation shown in red and negative correlation shown in light blue. The black dot represents the CpG shown in the scatter plot on the right. CGI represents a CpG island. K27 represents H3K27me3 in the matched normal tissue, shown as signal track (filled*

*polygon) shown above called peaks (box). K4 represents H3K4me3 in the matched normal tissue, shown as signal track (filled polygon) shown above called peaks (box). UCSC track represents gene transcripts obtained from UCSC. Right: Scatterplot of the DNA methylation (x-axis) and gene expression (y-axis) for the given CpG probe and the given gene. Each dot represents one sample (normal in green, tumour in purple). Spearman correlation coefficient R and p-value P are shown in the top left corner.*

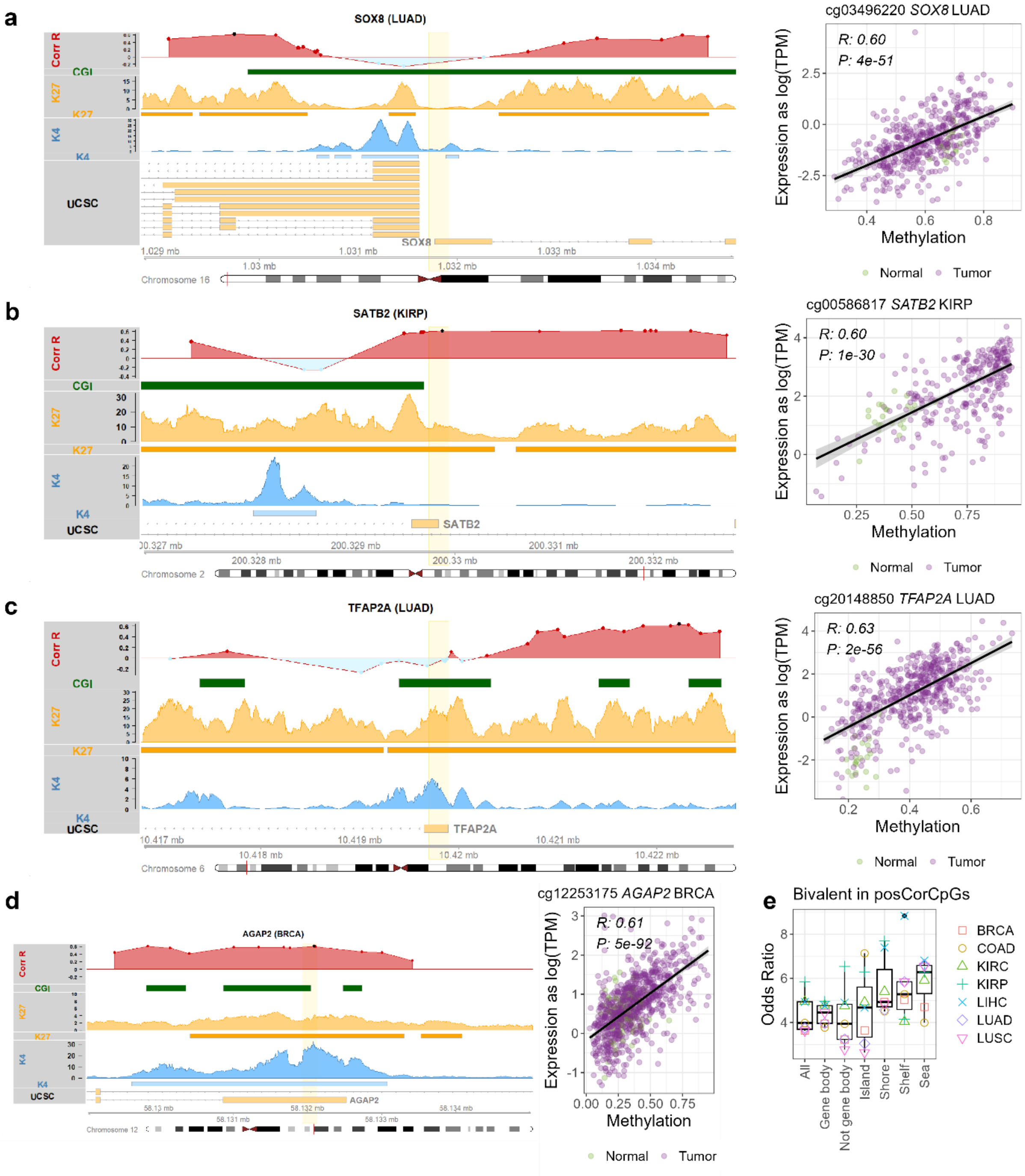

*Supplementary Figure 5:* ***Examples of bivalent genes with strongly positively correlated CpGs in their promoter regions. a-d,*** *Left: Genomic visualization of different (epi)genomic features in the example promoter regions: Corr R represents the Spearman Correlation coefficient between DNA methylation in the given CpG and*

*gene expression of the given gene, with positive correlation shown in red and negative correlation shown in light blue. The black dot represents the CpG shown in the scatter plot on the right. CGI represents a CpG island. K27 represents H3K27me3 in the matched normal tissue, shown as signal track (filled polygon) shown above called peaks (box). K4 represents H3K4me3 in the matched normal tissue, shown as signal track (filled polygon) shown above called peaks (box). UCSC track represents gene transcripts obtained from UCSC. Right: Scatterplot of the DNA methylation (x-axis) and gene expression (y-axis) for the given CpG probe and the given gene. Each dot represents one sample (normal in green, tumour in purple). Spearman correlation coefficient R and p-value P are shown in the top left corner.* ***e,*** *Positively correlated CpGs are enriched in bivalent promoters across different genomic categories. Odds ratio between bivalent promoters and positively correlated CpGs – shown across different categories of genomic regions. For example, the second boxplot shows enrichment of bivalent promoters and positively correlated CpGs, restricted only to CpGs located in gene bodies. It shows that positively correlated CpGs are enriched in bivalent promoters both when looking only at CpGs located inside gene bodies (second boxplot), as well as looking only outside gene bodies, so upstream TSS (third boxplot).*

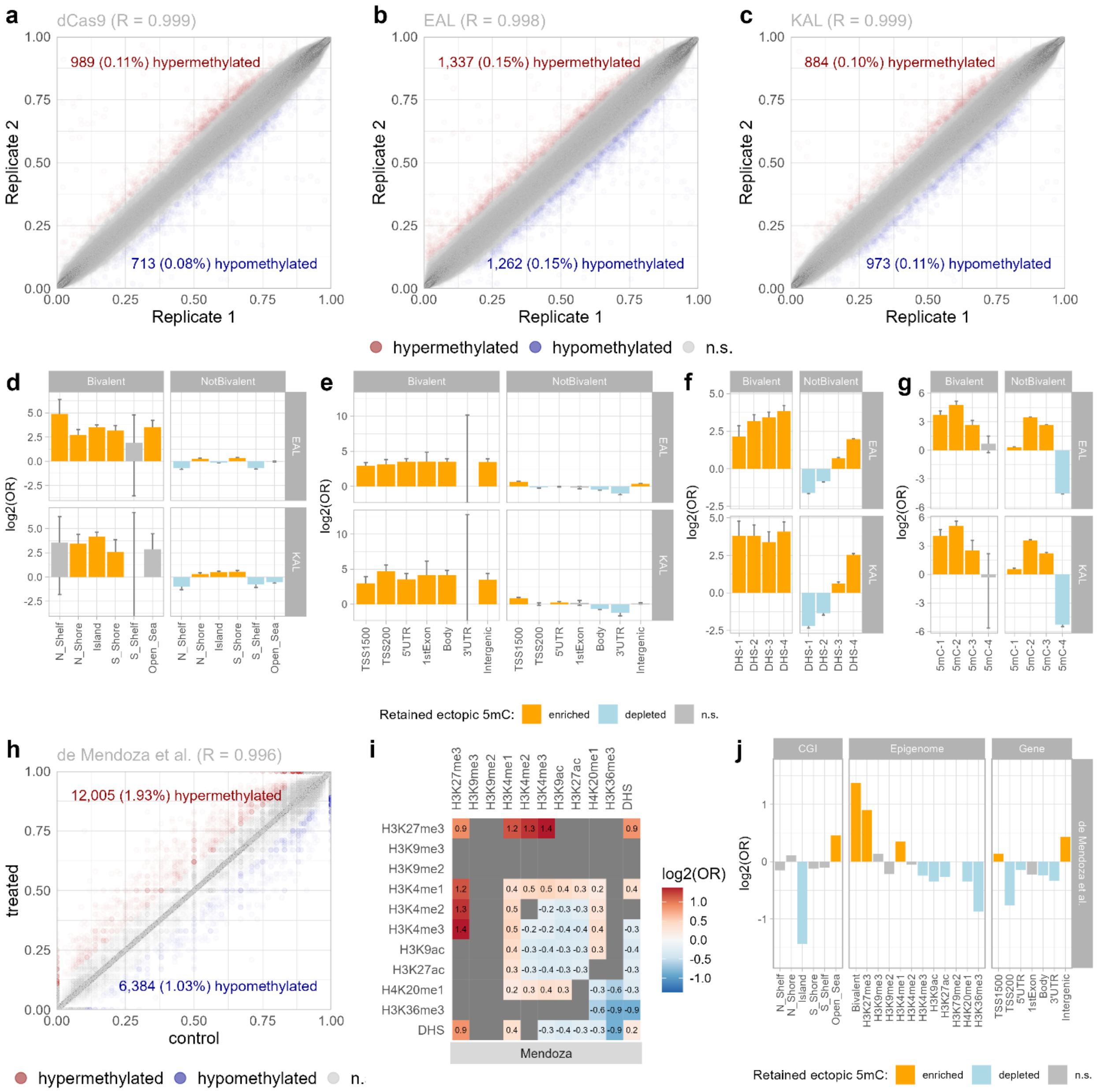

*Supplementary Figure 6:* ***Bivalent chromatin predisposes to long-lasting ectopic DNA methylation in CRISPR/dCas9 and Epi-ZF epigenome editing. a-c,*** *Scatterplot showing a correlation of EPIC array DNA methylation values between replicates. Each dot represents one CpG probe. The Pearson correlation coefficient R is shown on top.* ***d-g,*** *For each genomic category on the x-axis, we show the $\log_2$ odds ratio (OR) of the enrichment of retained methylation in regions where the (non)bivalent chromatin overlaps with the given genomic category. For example, the third bar in the Bivalent box represents enrichment of retained CpG methylation in bivalent regions overlapping with CpG islands. Orange represents positive enrichment, blue represents depletion, grey represents non-significant BH-adjusted p-value.* ***h,*** *DNA methylation values in treated and control samples in the Epi-ZF dataset by de Mendoza et al., 7 days post treatment withdrawal. The scatterplot shows the WGBS CpG methylation values in individual CpGs that are covered in the EPIC array. Each value represents combination of two replicates (see Methods). The Pearson correlation coefficient R is shown on top.* ***i,*** *Log2 odds ratio of the enrichment of retained methylated CpGs in genomic regions marked with a*

*combination of the given two histone marks in MCF7. Data from the Epi-ZF dataset by de Mendoza et al., 7 days post treatment withdrawal. Significant positive enrichment is shown in red, significant depletion is shown in blue, non-significant combinations are shown in grey. Benjamini-Hochberg (BH) adjusted Fisher's exact test p-value of at least 0.01 is considered significant.* ***j,*** *Enrichment of different genomic features and retained CpG methylation in the Epi-ZF dataset by de Mendoza et al., 7 days post treatment withdrawal. For each genomic category on the x-axis, we show the log2 odds ratio (OR) of enrichment of retained CpG methylation in the given genomic category. Orange represents positive enrichment, blue represents depletion, grey represents non-significant BH-adjusted p-value.*

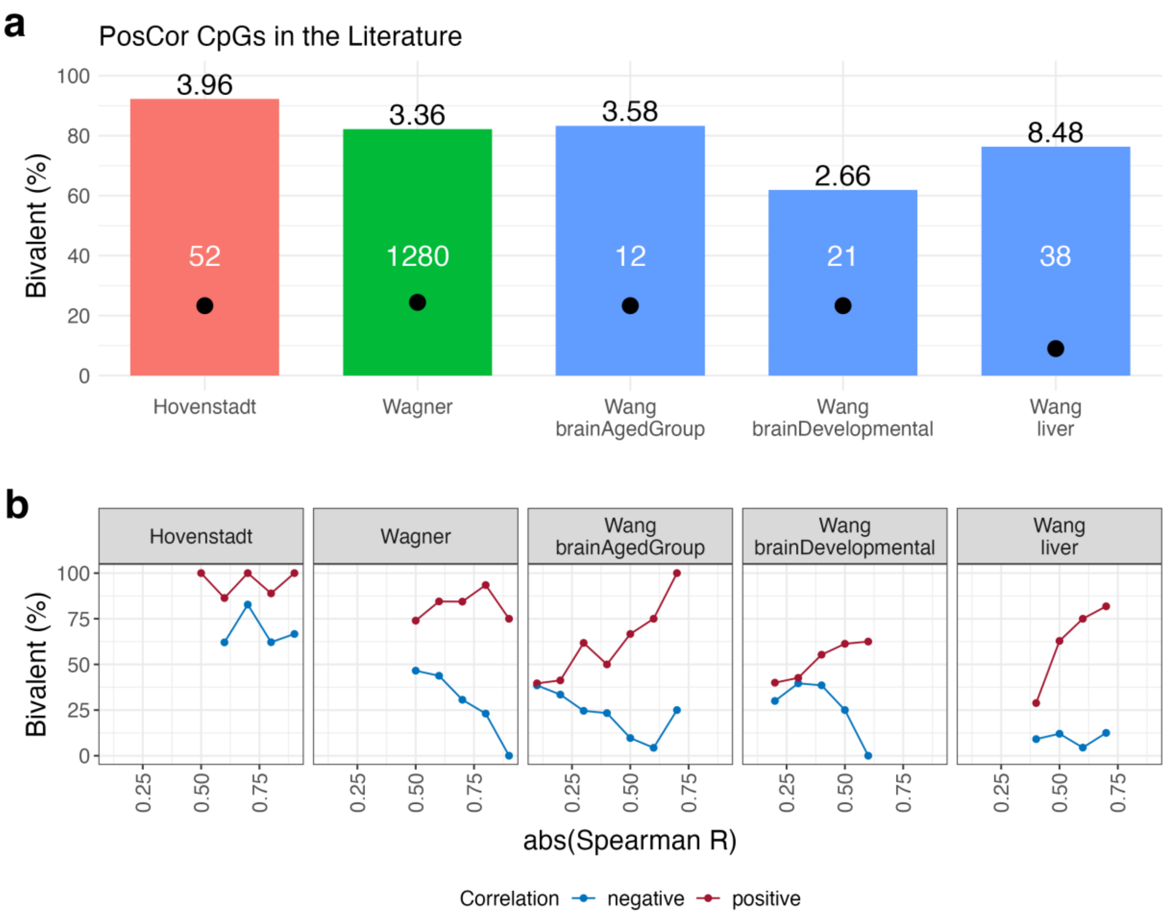

*Supplementary Figure 7:* ***Positively correlated promoter CpG-gene pairs in previously published studies are strongly enriched in bivalent chromatin.*** *Here, we analysed data from three studies that identified CpG-gene pairs with positive or negative correlation between DNA methylation and expression. These datasets include medulloblastoma (Hovenstadt et al.), primary fibroblasts from skin (Wagner et al.), and healthy tissues (Wang et al.) from ageing brain (25 to 78 years old), developing brain (foetus to age 25 years), and liver. Only groups with over 10 positively correlated CpGs were included. In all datasets, only rows with FDR < 0.05 were included.*

*In the Wang et al. dataset, only rows annotated as promoters were included. In the Hovenstadt et al. dataset, the rows represent genes, and each gene in the list is in vicinity of a positively correlated region within a DNA methylation valley. In the other two datasets, the rows represent CpG-gene pairs.*

*a, Percentage of positively correlated rows (correlation R ≥ 0.5) with bivalent promoters in matched normal tissue (data from ENCODE, see Supplementary Table 10). The total number of included rows is indicated in white. The expected percentage of bivalent rows is shown as a black dot, and computed based on genome-wide average. The numbers above the bars denote fold-change of observed over expected values.*

*b, Percentage of rows (CpG-gene pairs or genes) with bivalent histone marks in matched normal tissue, stratified by the value of the correlation R on the x-axis (positively correlated = red, negatively correlated = blue).*

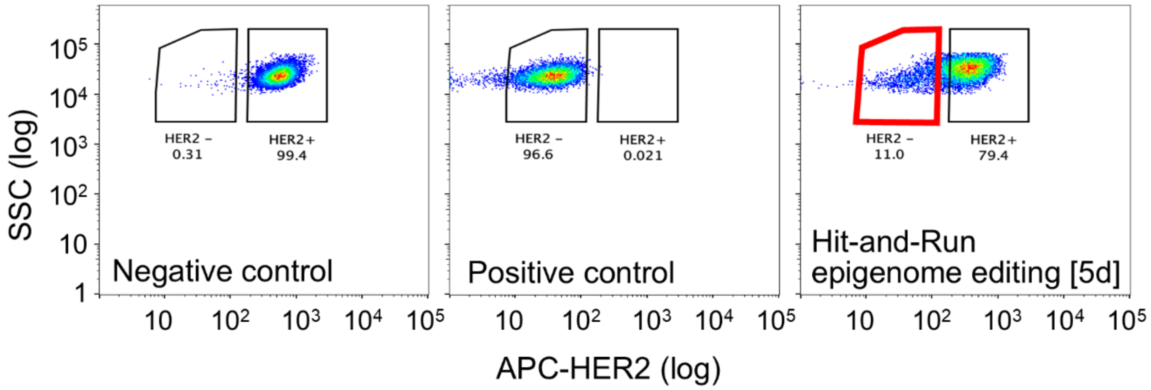

*Supplementary Figure 8:* ***Flow cytometry analysis of HER2-negative (HER2-) cells.*** *Representative FACS images show HER2 status using APC-HER2 antibody staining. HCT116 cells stained with APC-HER2 antibody (left) serve as a positive control (left). Unstained HCT116 cells (middle) serve as a negative control. A representative image (right) shows cells after transient transfection assays with KAL and gRNAs targeting HER2/ERBB2. HER2- cells, highlighted with a red gate, were collected 5 days after hit-and-run epigenome editing. Plots display side scatter (SSC) on the y-axis and forward scatter FSC (640-671nm), measuring APC-HER2, on the x-axis.*
